# Inosine is an alternative carbon supply that supports effector T cell proliferation and antitumor function under glucose restriction

**DOI:** 10.1101/766642

**Authors:** Tingting Wang, JN Rashida Gnanaprakasam, Xuyong Chen, Siwen Kang, Xuequn Xu, Hua Sun, Lingling Liu, Ethan Miller, Teresa A. Cassel, Qiushi Sun, Sara Vicente-Muñoz, Marc O. Warmoes, Andrew N. Lane, Xiaotong Song, Teresa W.-M. Fan, Ruoning Wang

## Abstract

T cells undergo a characteristic metabolic rewiring that fulfills the dramatically increased bioenergetic, biosynthetic, and redox demands following antigen stimulation. A robust adaptive immune system requires effector T cells to respond and adapt to fluctuations in environmental nutrient levels imposed by infectious and inflammatory sites in different tissues. Inevitably, such responsiveness and adaptation reflect metabolic plasticity, allowing T cells to elicit immune functions by using a wide range of nutrient substrates. Here, we show that effector T cells utilize inosine, as an alternative substrate, to support cell growth and function in the absence of glucose. T cells metabolize inosine into hypoxanthine and phosphorylated ribose by purine nucleoside phosphorylase (PNP). Using Stable Isotope-Resolved Metabolomics (SIRM), we demonstrated that ribose moiety of inosine can enter into central metabolic pathways to provide ATP and biosynthetic precursors. Accordingly, the dependence of T cells on extracellular glucose for growth and effector functions can be relieved by inosine. On the other hand, cancer cells display diverse capacity to utilize inosine as a carbon resource. Moreover, the supplement of inosine enhances the anti-tumor efficacy of immune-checkpoint blockade or adoptive T cell transfer, reflecting the capability of inosine in relieving tumor-imposed metabolic restrictions on T cells *in vivo*.

## Introduction

A robust adaptive immune response relies on the ability of antigen-specific T cells to rapidly transform from a quiescent (naive) state to a proliferative (active) state, followed by sustained proliferation during the period of antigen presentation. An increasing body of evidence suggests that a coordinated rewiring of cellular metabolism, is required to fulfill the bioenergetic, biosynthetic and redox demands of T cells following activation [1; 2; 3; 4; 5; 6; 7; 8; 9; 10; 11]. Naive T cells and memory T cells predominantly rely on fatty acid oxidation (FAO) and oxidative phosphorylation (OXPHOS) for their energy supply in the quiescent state. Following antigen stimulation, effector T (T_eff_) cells rapidly upregulate other pathways including aerobic glycolysis, the pentose phosphate pathway (PPP) and glutaminolysis to drive clonal expansion and effector functions [12; 13; 14; 15; 16; 17; 18; 19; 20]. Such metabolic reprogramming induced by T cell activation relies on a hierarchical signaling cascade transcriptional networks [13; 16; 21; 22; 23; 24; 25; 26].

It is now commonly recognized that the immune system is intimately involved in tumor initiation, progression, and responses to therapy [27; 28; 29; 30; 31; 32]. T_eff_ cells are the major agents of antitumor immunity, and elicit anti-tumor activity through direct recognition and killing of antigen presenting tumor cells as well as orchestrating a plethora of adaptive and innate immune responses [31; 33; 34; 35]. However, tumors often co-opt a broad spectrum of mechanisms that foster an immune suppressive microenvironment, thereby escaping T cell-mediated anti-tumor immune response. Various intrinsic and extrinsic tumor factors favor the development of immunosuppressive myeloid-derived suppressor cells (MDSC) and regulatory T (T_reg_) cells, increasing the expression of inhibitory checkpoint receptors, such as cytotoxic T-lymphocyte-associated protein 4 (CTLA-4), programmed cell death protein 1 (PD-1), and lymphocyte-activation gene 3 (LAG 3); while reducing the expression and presentation of tumor specific antigens [30; 36; 37; 38; 39; 40; 41]

Mounting evidence has shown that immunotherapies through strengthening the amplitude and quality of T cell-mediated adaptive response may mediate durable and even complete tumor regression in some cancer patients. Two of the most promising approaches to enhance therapeutic anti-tumor immunity are immune checkpoint blockade using monoclonal antibodies against PD-1/PDL1 and CTLA4, and adoptive cell transfer (ACT) of tumor infiltrating lymphocytes (TILs) or peripheral T cells that are genetically engineered with chimeric antigen receptors (CARs) [30; 42; 43; 44; 45; 46; 47; 48; 49]. Despite the clinical promise of CAR T-cell therapy in B cell leukemia and checkpoint blockade therapies in metastatic melanoma, non-small cell lung cancer (NSCLC), bladder cancer, and others, more than three-quarters of cancer patients overall remain refractory to the current immunotherapeutic regimen [49; 50; 51; 52; 53; 54].

To enhance immunotherapeutic efficacy, understanding the key immunosuppressive barriers in cancer tissues is critically important. A key but largely overlooked problem is nutrient competition between tumor cells and T cells in the nutrient poor tumor microenvironment (TME). Metabolic dysregulation is now recognized as one of the hallmarks of human cancer and the activation of oncogenic signaling pathways enabling tumor cells to reprogram pathways of nutrient acquisition and metabolism to meet the additional bioenergetic, biosynthetic and redox demands of cell transformation and proliferation. [55; 56; 57; 58; 59; 60; 61; 62; 63]. The TME of solid tumors represents a dramatic example of metabolic stress, where the high metabolic demands of cancer cells can restrict the function of T_eff_ cells by competing for nutrients including glucose and by producing immune suppressive metabolites [62; 64; 65; 66; 67; 68; 69; 70; 71; 72; 73; 74]. As such, a better understanding of the metabolic modulation of T_eff_ cells to relieve the immunosuppressive barriers in the TME will enable us to devise rational and effective approaches to enhance cancer immunotherapy via improving T cell metabolic fitness.

Here, we report that T cells can utilize few substrates, and that inosine is an alternative metabolic substrate that can support cell growth and crucial T-cell functions in the absence of glucose. The catabolism of the ribose subunit of inosine provides both metabolic energy in the form of ATP and biosynthetic precursors from glycolysis and the pentose phosphate pathway. Inosine supplementation promotes T cell-mediated tumor killing activity *in vitro* and enhances the antitumor efficacy of checkpoint blockade therapy or adoptive T cell therapy in mouse models, reflecting the capability of inosine in relieving tumor-imposed metabolic restrictions on T cells.

## Results

### A metabolic screen identifies inosine as an alternative fuel to support T_eff_ cell survival and proliferation

Solid tumors typically develop a hostile microenvironment characterized by an irregular vascular network and a correspondingly poor nutrient supply. Highly glycolytic cancer cells may deplete nutrients, particularly glucose, thereby restricting glucose availability to T cells [66; 67; 68; 72]. Previous studies have shown that murine T cells significantly enhanced glucose catabolic activity following activation, and that glucose starvation compromises viability and proliferation of T_eff_ cells [13; 14; 16; 25; 75]. We asked whether T_eff_ cells have the capability of taking up alternative carbon and energy sources to support their survival and proliferation in the absence of glucose. To this end, we generated human T_eff_ cells by stimulating human PBMCs with plate-bound anti-CD3 antibody in the presence of IL-2. After three days activation and expansion of human T_eff_ cells, we plated cells in glucose-free media into 96 well plates (Biolog’s Phenotype MicroArray Mammalian plates PM-M1 and PM-M2) pre-loaded with an array of compounds that can serve as carbon and/or nitrogen sources for mammalian cells. Blank wells and wells pre-loaded with glucose were included as negative and positive controls, respectively. We then determined the substrate utilization of T_eff_ cells in each well through a colorimetric assay in which the formation of the reduced dye represents cells’ ability to catabolize these extracellular substrates and generate NADH [76]. The results were normalized to positive control (wells preloaded with glucose) and summarized in Figure 1A and Figure S1A. It was found that polysaccharides, six-carbon sugars and their derivatives including dextrin, glycogen, maltotriose, maltose, mannose and fructose-6-phosphate were utilized by T_eff_ cells in the absence of glucose (Figure 1A and Figure S1A). Remarkably, inosine, a nucleoside, also supported T_eff_ cells bioenergetic activity in the absence of glucose (Figure 1A). To eliminate the possibility that inosine was contaminated with glucose or glucose analogs, we obtained inosine from a different source and determined if inosine could support mouse and human T_eff_ cell proliferation and viability in the absence of glucose. Freshly isolated mouse and human CD8+ T cells were stained with carboxyfluorescein succinimidyl ester (CFSE) dye and stimulated with plate-bound anti-CD3 and anti-CD28 antibodies in conditional media supplied with interleukin 2 (IL-2). Glucose starvation significantly reduced cell proliferation and increased cell death (Figure 1B, 1C, S1B and S1C). However, Inosine supplemented at an equimolar amount of glucose markedly reduced cell death and restored cell proliferation in mouse and human T_eff_ cells following glucose starvation (Figure 1B, 1C, S1B and S1C). This shows that inosine may offer bioenergetic support similar to as glucose to support T_eff_ cell proliferation.

**Figure 1.**
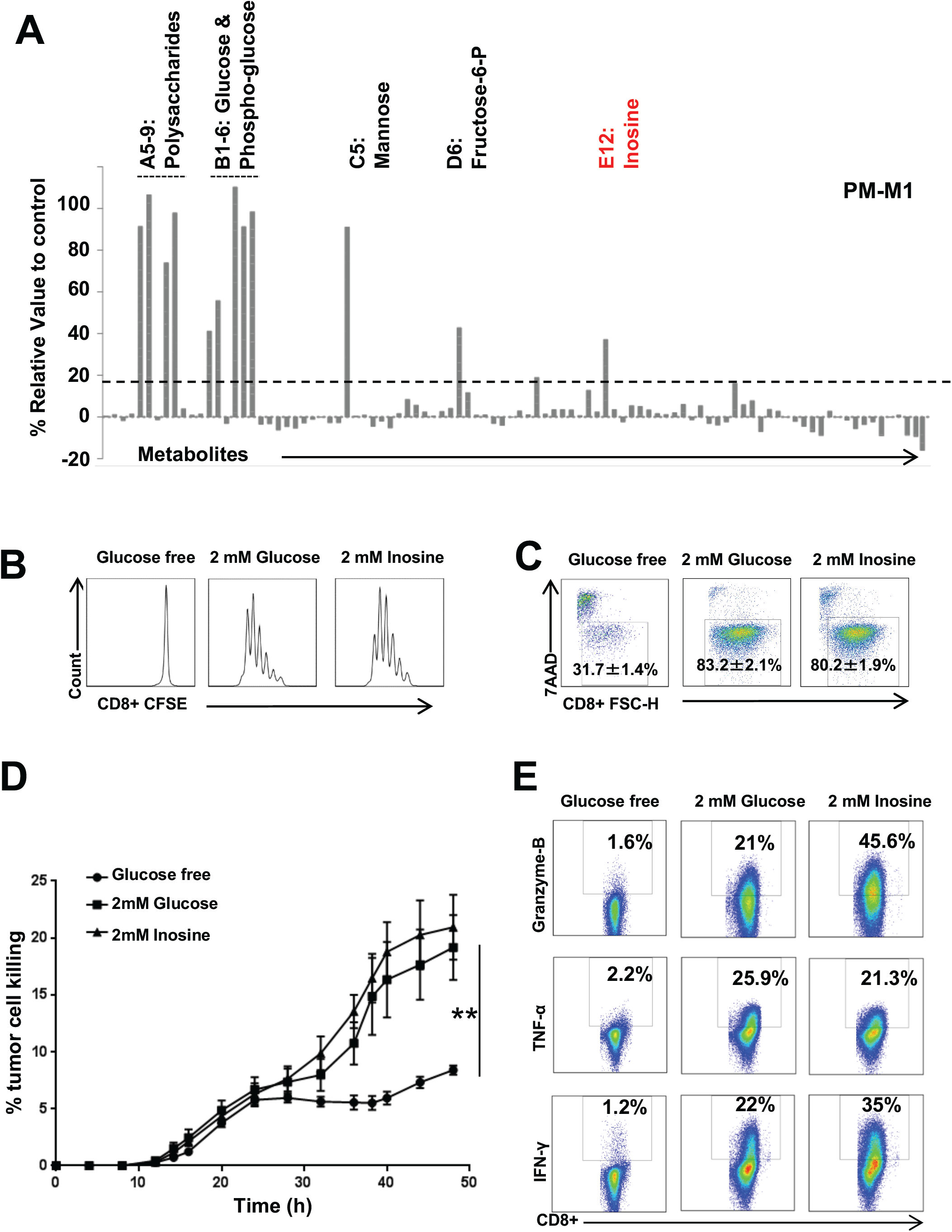
Inosine can support T effector cells proliferation and function in the absence of glucose. (A) Metabolite array from the PM-M1 Microplate (Biolog). Activated human T cells were incubated with various energy sources in a PM-M1 Microplate for 24 h, followed by Biolog redox dye mix MB incubation and measured spectrophotometrically at 590 nm. (B-C) Naive CD8+ T cells from C57BL/6 mice were activated by plate-bound anti-CD3 and anti-CD28 antibodies in complete media for 24 h, then the cells were switched to indicated conditional media and were cultured for 48 h. Cell proliferation and cell death were determined by CFSE dilution and 7AAD uptake, respectively. Data are representative of three independent experiments. (D) B16F10 tumor cells were co-cultured with activated Pmel T cells in the indicated conditional media, and tumor cell death was evaluated using caspase 3/7 reagent by IncuCyte ZOOM™. Error bars represent mean ± SD (n=4). **, *P*=0.0072 for glucose free versus 2 mM Inosine. (E) Naive CD8^+^ T cells from C57BL/6 mice were activated by plate bound anti-CD3 and anti-CD28 antibodies and differentiated in the indicated conditional media for 4 days. The indicated proteins were quantified by intracellular staining following PMA and ionomycin stimulation.

### Inosine supports T effector cell function in the absence of glucose

We next asked whether inosine could restore the cytotoxic activity and other effector functions of T_eff_ cells in the absence of glucose. We generated antigen specific mouse T_eff_ cells by stimulating major histocompatibility complex (MHC) class I-restricted T cells, isolated from premelanosome protein (Pmel)-1 T cell receptor transgenic mice, with peptide antigen and interleukin 2 (IL-2) [77; 78; 79]. We also generated human GD2-specific chimeric antigen receptor (GD2-CAR) T cells by stimulating human PBMCs with plate-bound anti-CD3 antibody in the presence of IL-2, followed by retroviral transduction with GD2-CAR [80; 81]. It was found that Pmel+ cells recognized Pmel-17 (mouse homologue of human SIVL/gp100) in mouse melanoma, while GD2-CAR T cells recognized GD2, a disialoganglioside expressed in tumors of neuroectodermal origin. We co-cultured Pmel+ T_eff_ cells with B16F10 mouse melanoma cells (Figure 1D), as well as GD2-CAR T cells with LAN-1 human neuroblastoma cells (Figure S1D), in conditional media, then assessed tumor cell apoptosis by real-time imaging analysis of caspase activity. We also cultured mouse (Figure 1E) and human (Figure S1E) T_eff_ cells in conditional media and assessed the expression of effector molecules including granzyme B, tumor necrosis factor alpha (TNF-α) and interferon gamma (IFNγ), all of which are important in mediating tumor cell-killing activity of T_eff_ cells. While glucose starvation significantly reduced the tumor-killing capacity and the expression of granzyme B, TNF-α and IFNγ of T_eff_ cells, both glucose and inosine supplementation restored these properties in mouse and human T_eff_ cells (Figure 1D, 1E, S1D and S1E). Taken together, our results indicate that inosine has the capacity to replace glucose in supporting the effector function of T_eff_ cells.

### Adenosine, the metabolic precursor of inosine, cannot support the proliferation and function of T effector cells in the absence of glucose

Adenosine, the metabolic precursor of inosine, has been reported as an immune suppressive metabolite and promotes the resolution of inflammation by targeting a range of immune cells, including T cells [82; 83; 84; 85]. Since adenosine and inosine have identical chemical properties, we asked whether adenosine can support the proliferation of T_eff_ cells in the absence of glucose. Adenosine supplemented at an equimolar amount of glucose, failed to reduce cell death or restore the proliferation of mouse and human T_eff_ cells following glucose starvation (Figure S2A and S2B). Next, we asked whether adenosine can support the effector functions of T_eff_ cells in the absence of glucose. We found that, while adenosine had a marginal effect of enhancing the expression of granzyme B, it failed to restore the tumor-killing activity and the expression of other effector molecules in mouse (Figure S2C and S2D) or human T_eff_ cells (Figure S2E and S2F) in the absence of glucose. Collectively, our results indicate that inosine, but not its metabolic precursor adenosine, has the capacity for replacing glucose in supporting the cytotoxic function of T_eff_ cells.

### Inosine does not support the proliferation of T effector cells by enhancing glutamine and fatty acid catabolism or through purine nucleotide salvage

Glutamine is a key carbon and nitrogen donor for T_eff_ cells, while fatty acids are also known carbon donors for naive T (Tn_ai_), regulatory T (T_reg_) and memory T (T_mem_) cells [5; 6; 11; 86]. We next asked whether inosine supports T_eff_ cell proliferation by enhancing glutaminolysis and fatty acid β-oxidation (FAO). Mouse T_eff_ cells cultured in glucose or inosine conditional media displayed comparable catabolic activities via glutaminolysis, as indicated by ^14^CO_2_ release from [^14^C_5_]-glutamine, or via FAO, indicated by ^3^H-water release from [9,10^−3^H]-palmitic acid (Figure. S3A). Moreover, an inhibitor of FAO (Etomoxir), failed to block T_eff_ cell proliferation in inosine conditional media (Figure. S3B). However, glutamine starvation blocks inosine-dependent T_eff_ cell proliferation (Figure. S3B), supporting the indispensable role of glutamine as a key nitrogen donor of T_eff_ cells [5; 6]. Collectively, our results show that inosine does not enhance glutamine and fatty acid catabolism in T_eff_ cells. Inosine is a hypoxanthine nucleoside that can be broken down into ribose-1-phosphate and hypoxanthine, the latter of which can funnel into purine nucleotide salvage pathway (Figure S3C) [87; 88]. We next asked whether hypoxanthine or the purine nucleoside guanosine can support the proliferation of T_eff_ cells in the absence of glucose. Neither hypoxanthine nor guanosine supplemented at an equimolar amount of glucose can reduce cell death or restore the proliferation of mouse T_eff_ cells following glucose starvation (Figure S3D). Together with our adenosine data, we conclude that purine nucleotide salvage is not responsible for inosine-dependent T_eff_ cell proliferation.

### Inosine-ribose fuels key metabolic pathways in T effector cells

Next, we asked whether inosine-derived ribose (Figure S3C) could be a fuel for key metabolic pathways involved in cell proliferation and survival. To this end, we applied a stable isotope based metabolic flux approach to compare metabolic routes of glucose and the ribose moiety in inosine. Specifically, we supplied [^13^C_6_]-Glucose (Glc) or [1’,2’,3’,4’,5’-^13^C_5_]-Inosine (Ino), where only the ribose contains carbon-13 (^13^C), as the metabolic tracer in human T_eff_ cells cultured with glucose free conditional media (Figure 2A). Then, we followed ^13^C incorporation into intermediate metabolites in the central carbon metabolic pathways including the pentose phosphate pathway (PPP), glycolysis and the Krebs cycle.

**Figure 2.**
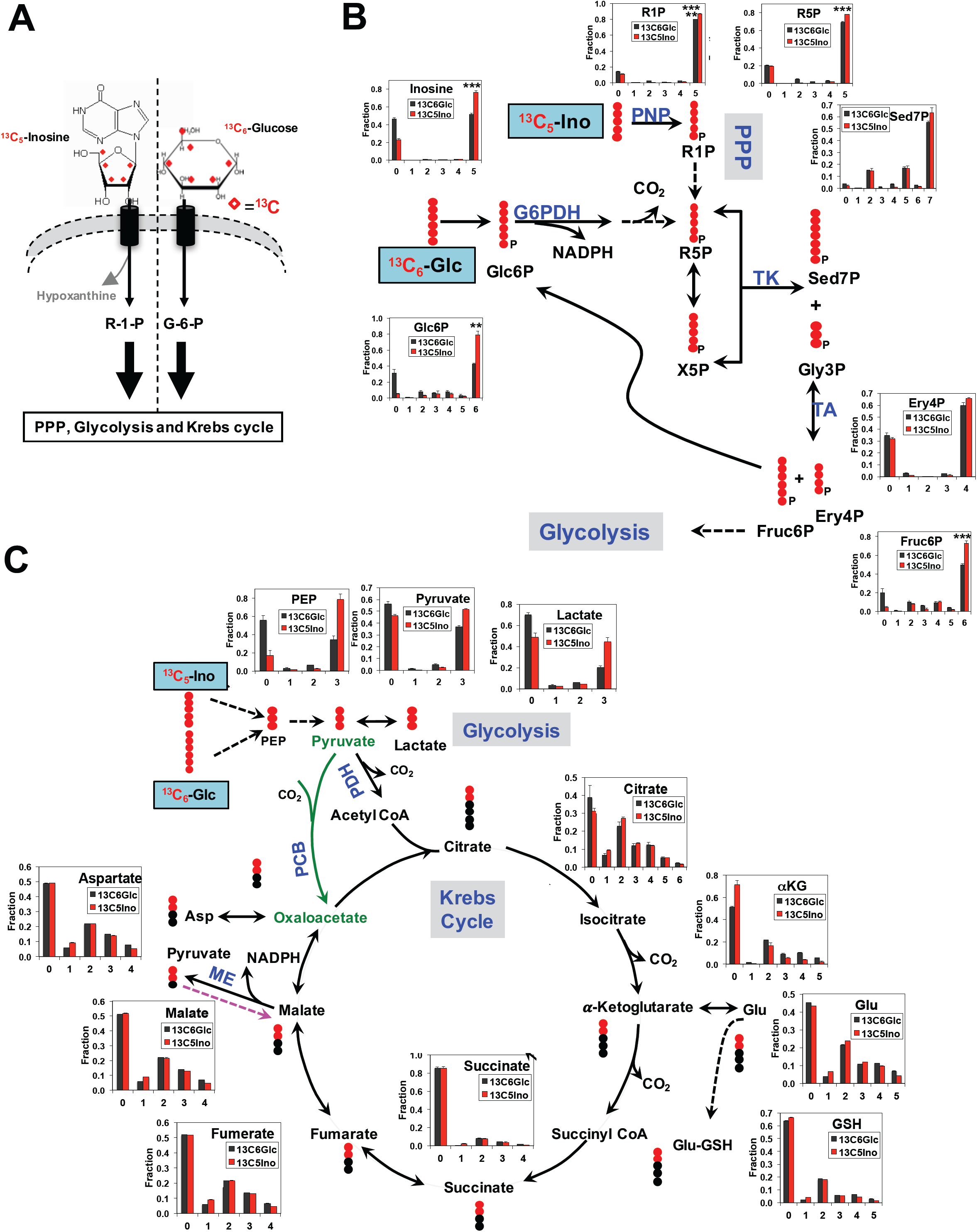
The ribose moiety of inosine can replace glucose and feed into the central carbon metabolism in T effector cells. (A) Diagram of [^13^C_5_]-inosine or [^13^C_6_]-Glucose uptake and conversion to ribose-1-phosphate (R-1-P) or glucose-6-phosphate (G-6-P), respectively, which subsequently enters the downstream PPP, glycolysis and Krebs cycle metabolic pathways. denoted the uniformly ^13^C label positions of the ribose moiety of inosine and all carbons of glucose. (B-C) Active human T cells were incubated in [^13^C_6_]-Glucose (Glc) or [^13^C_5_]-ribose-inosine (Ino) for 24 h, extracted as described in methods, and analyzed for PPP metabolites (B) and glycolytic/Krebs cycle metabolites (C) by IC-UHR-FTMS. Glc6P: glucose-6-phosphate; R1P: ribose-1-phosphate; R5P: ribose-5-phosphate; X5P: xylulose-5-phosphate; Sed7P: sedoheptulose-7-phosphate; Gly3P: glyceraldehyde-3-phosphate; Ery4P: erythrose-4-phosphate; G6PDH: glucose-6-phosphate dehydrogenase; TK: transketolase; TA: transaldolase; Fruc6P: fructose-6-phosphate; *α*KG: *α–*ketoglutarate; Glu-GSH: glutamyl unit of glutathione; PDH: pyruvate dehydrogenase; PCB: pyruvate carboxylase; ME: malic enzyme; •: ^12^C; •: ^13^C derived either from the PDH or PCB-initiated Krebs cycle reactions, or from the reverse ME reaction. Solid and dashed arrows represent single- and multi-step reactions, single and double-headed arrows refer to irreversible and reversible reactions, respectively. Numbers in the X-axis represent those of ^13^C atoms in given metabolites.

We first compared the metabolism of [^13^C_6_]-Glc versus [^13^C_5_]-Ino via the PPP (Figure 2B). As expected, the ^13^C_5_ fractional enrichment of the parent tracer inosine were higher in the inosine tracer-treated T_eff_ cells than the glucose tracer-treated ones. Moreover, both tracers were extensively metabolized via the PPP to produce ^13^C labeled intermediates, including ribose-5-phosphate (R1P), ribose-5-phosphate (R5P), sedoheptulose-7-phosphate (Sed7P), erythrose-4-phosphate (Ery4P), fructose-6-phosphate (Fruc6P) and glucose-6-phosphate (Glc6P) (Figure 2B). The fraction of ^13^C enrichment in the dominant fully ^13^C labeled isotopologues of R5P, Fruc6P and Glc6P were higher for the inosine tracer treatment than the glucose tracer treatment. This is consistent with the direct and rapid metabolism of [^13^C_5_]-inosine via the non-oxidative branch of the PPP and the equilibration of Fruc6P with Glc6P via the action of phosphoglucose isomerase (PGI). We also observed the presence of scrambled ^13^C products of Sed7P ([^13^C_2_]- and [^13^C_5_]-Sed7P in particular), Fruc6P, and Glc6P. These scrambled products were presumably derived from the reversible transketolase (TK) and transaldolase (TA) activity. The dominant presence of [^13^C_2_]- and [^13^C_5_]-isotopologues among the ^13^C scrambled products of Sed7P is again consistent with the flux through both branches of PPP for both inosine and glucose tracer treatments.

The ^13^C-Fruc6P and Glc6P products of [^13^C_5_]-inosine from PPP can be further metabolized via glycolysis and the Krebs cycle, as in the case of [^13^C_6_]-Glc. We thus tracked the fate of [^13^C_5_]-inosine and [^13^C_6_]-Glc in human T_eff_ cells through these two key bioenergetic pathways via IC-UHR-FTMS analysis (Figure 2C). [^13^C_5_]-inosine was metabolized as extensively as [^13^C_6_]-Glc via glycolysis to produce fully ^13^C labeled isotopologues of intermediate metabolites including phosphoenolpyruvate (PEP), pyruvate, and lactate. The presence of a significant fraction of [^13^C_2_]-isotopologues among the ^13^C products of the Krebs cycle (citrate, *α–*ketoglutarate or *α*KG, glutamate or Glu, glutathione or GSH, succinate, fumarate, malate, and aspartate or Asp) is consistent with flux through pyruvate dehydrogenase (PDH)-dependent reaction for both inosine and glucose tracer treatments (Figure 2C). Together, these data support an extensive capacity of T_eff_ cells to utilize inosine as an alternative fuel source to glucose in the central carbon metabolism.

### The Inosine hydrolyzing enzyme, purine nucleoside phosphorylase, is required for inosine-dependent proliferation and effector functions in T effector cells

Purine nucleoside phosphorylase (PNP) is responsible for hydrolyzing inosine into ribose-1-phosphate and is significantly induced following T cell activation (Figure 3A and 3B) [87; 88]. Given that the ribose moiety of inosine is extensively catabolized in T_eff_ cells, we reasoned that PNP is required for inosine-mediated bioenergetic support. To test this idea, we assessed the bioenergetic activity of T_eff_ cells in the presence of a PNP inhibitor (forodesine (Foro); also referred as BCX-1777, Immucillin H) [89; 90]. As shown in Figure 3C and S4A, the PNP inhibitor Foro treatment led to a dosage-dependent attenuation of bioenergetic activity in both human and mouse T_eff_ cells cultured in inosine-containing but not glucose-containing media. We then assessed the proliferation, viability and effector functions of T_eff_ cells treated with Foro. Consistent with its effects on the bioenergetic activity, Foro significantly dampened proliferation, viability, tumor killing activity as well as the expression of granzyme B, TNF-α and IFNγ in mouse (Figure 3D-3F) and human (Figure S4B-S4D) T_eff_ cells cultured in inosine-containing media. Next, we sought to validate the data of pharmacological inhibition of PNP by knocking down PNP in human T_eff_ cells. As shown in Figure S4E and S4F, electroporation with PNP siRNA but not control scramble siRNA partially reduced the level of PNP protein in T_eff_ cells, and attenuated the bioenergetic activity and number of human T_eff_ cells cultured in inosine-containing but not glucose-containing media. Taken together, our results suggest that PNP-dependent inosine hydrolysis is required for inosine-mediated proliferation, bioenergetic support, tumor killing activity, and the expression of T_eff_ cell effector molecules.

**Figure 3.**
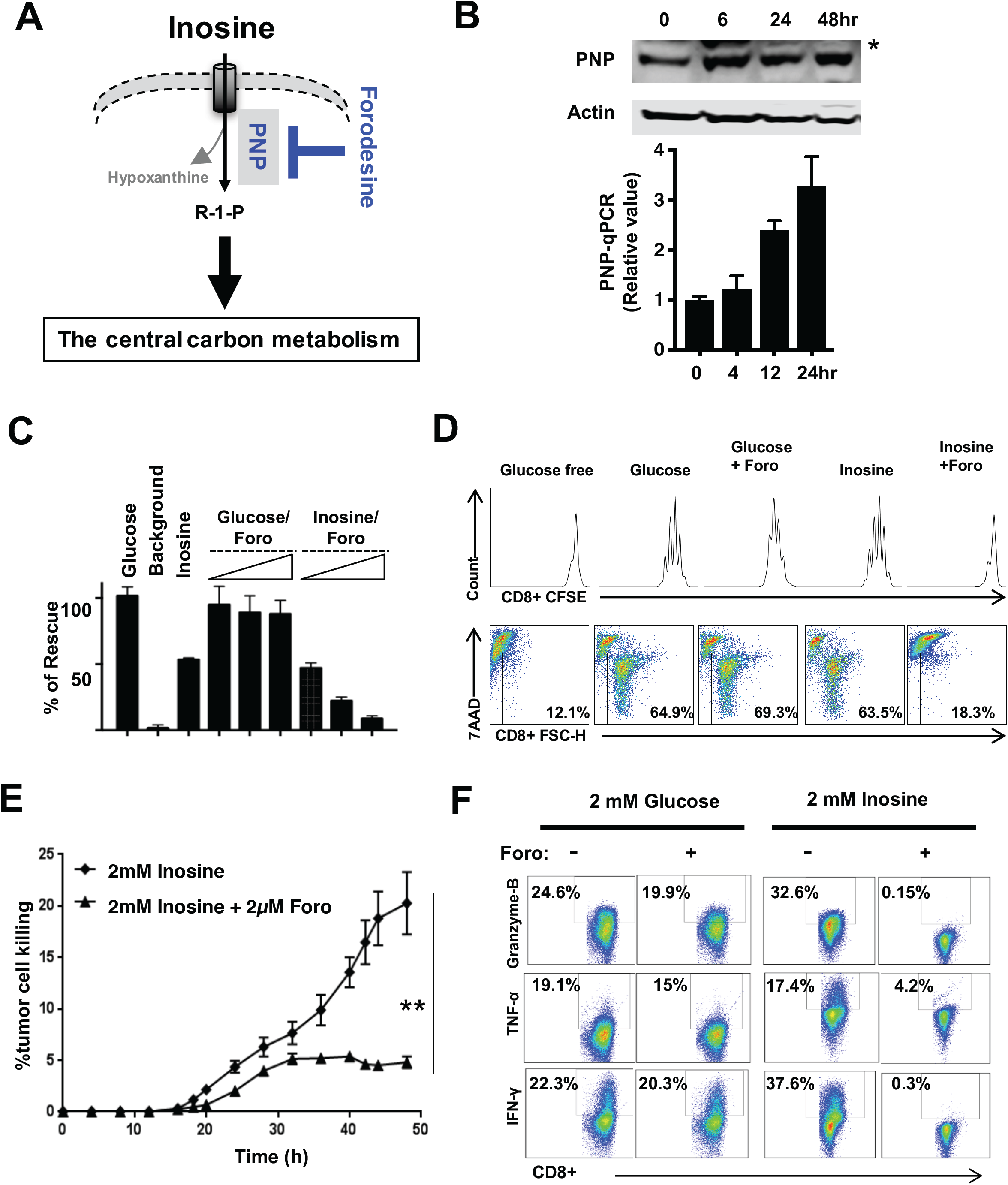
The purine nucleoside phosphorylase (PNP) is required for inosine-dependent mouse T effector cell proliferation and effector functions. (A) Diagram of purine nucleoside phosphorylase (PNP) inhibitor Foroesine (Foro) in blocking the breakdown of inosine into hypoxanthine and R-1-P. (B) PNP protein and mRNA expression levels in mouse T cells at the indicated time points upon activation by plate-bound anti-CD3 and anti-CD28 antibodies were determined by immunoblot (upper) and qPCR (lower), respectively. * indicates a nonspecific band recognized by PNP antibody. Error bars represent standard deviation from the mean of triplicate samples. Data are representative of two independent experiments. (C) Activated mouse T cells were incubated without glucose (background), with glucose or with inosine, as well as in combination with increased concentrations of Foro (0.1 μM, 0.5 μM, and 2 μM) for 24 h, followed by Biolog redox dye mix MB incubation and measured spectrophotometrically at 590 nm. (D) Naive CD8+ cells from C57BL/6 mice were activated by plate-bound anti-CD3 and anti-CD28 antibodies in complete media for 24 h, then the cells were switched to the indicated conditional media in combination with 2 μM Foro for 72 h. Cell proliferation and cell death were determined by CFSE dilution (upper) and 7AAD uptake (lower), respectively. Data are representative of three independent experiments. (E) B16F10 tumor cells were co-cultured with activated Pmel T cells in the presence of inosine with or without Foro, and tumor cell death was evaluated using caspase 3/7 reagent by IncuCyte ZOOM™. Error bars represent mean ± SD (n=4). **, *P*<0.01 for 2 mM Inosine versus 2 mM Inosine + 2 μM Foro. Data are representative of two independent experiments. (F) Naive CD8+ T cells from C57BL/6 mice were activated by plate bound anti-CD3 and anti-CD28 antibodies and differentiated in the indicated conditional media with or without 2 μM Foro for 4 days. The indicated proteins were quantified by intracellular staining following PMA and ionomycin stimulation. Data are representative of three independent experiments.

### The PNP inhibitor, Forodesine, suppresses inosine catabolism in T effector cells

Next, we assessed the effect of PNP inhibition on inosine catabolism by comparing the carbon flux of [^13^C_5_]-Inosine in the presence or absence of the PNP inhibitor (Foro), in T_eff_ cells (Figure 4A). Since Foro treatment significantly reduces T cell viability in inosine-containing media (Figure 3D and S4B), we chose to moderately suppress the PNP activity by applying a lower dose of Foro along with a shorter incubation time compared to the previous experiment. Treatment with Foro led to the fractional enrichment of ^13^C_5_ inosine, the metabolic substrate of PNP, and a decrease in the fractional enrichment of the fully ^13^C labeled species of PPP metabolites, which are indirect metabolic products of PNP (Figure 4B). These results suggested that the inhibition of PNP suppresses the catabolism of the ribose moiety of inosine via the PPP. Next, we tracked the fate of [^13^C_5_]-inosine via glycolysis and the Krebs cycle in the context of suppressed PNP activity. As shown by IC-UHR-FTMS analysis (Figure 4C), the Foro treatment greatly reduced the fraction of fully ^13^C labeled isotopologues of glycolysis metabolites and the fraction of [^13^C_2_]-isotopologues of the Krebs cycle metabolites. To further assess the effect of Foro in T cell metabolism in both glucose- and inosine-containing media, we applied 1D HSQC NMR to quantify ^13^C labeled isotopologues of several representative metabolites in glycolysis, the Krebs cycle and nucleotide biosynthesis, including lactate (Lac), alanine (Ala), inosine (Ino), glutamine (Glu), GSH+GSSG (Gluta), adeninine nucleotides (AXP) and uracil nuecleotides (UXP) (Figure S5). Foro treatment significantly altered the quantity of [^13^C_5_]-Ino-derived metabolites (Figure S5A). In contrast, Foro had little effect on the quantity of [^13^C_6_]-Glc-derived metabolites (Figure S5B). Together, these data support an extensive capacity of T_eff_ cells for hydrolyzing inosine as an alternative fuel source to glucose in the central carbon metabolism.

**Figure 4.**
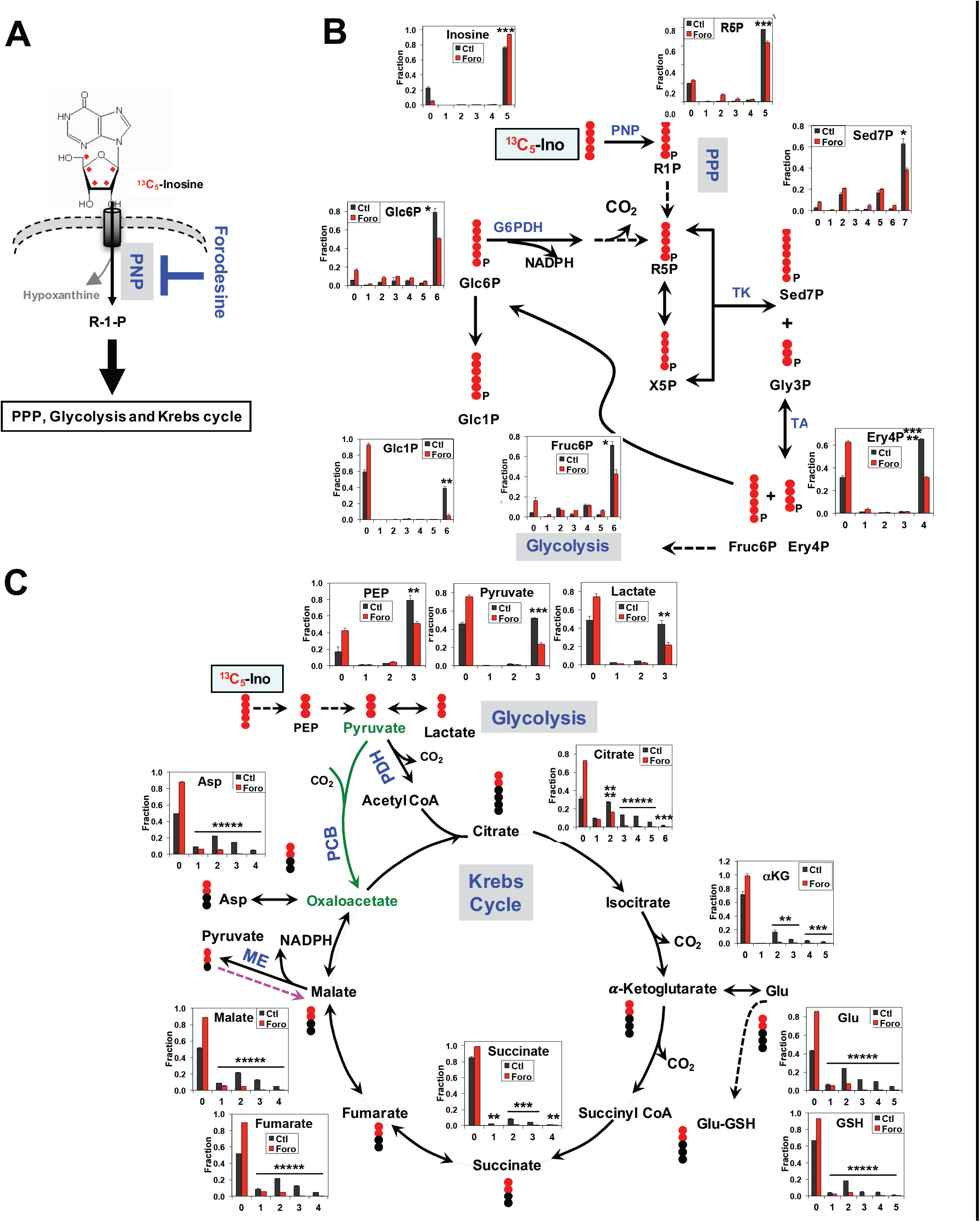
PNP inhibitor forodesine suppresses inosine catabolism in T effector cells. (A) Diagram of PNP inhibitor Forodesine (Foro) blocking the catabolism of [^13^C_5_]-inosine into R-1-P and its subsequent entrance into the PPP, Glycolysis and Krebs cycle. ♦ denoted the uniformly ^13^C label positions of the ribose moiety of inosine. (B-C) Active human T cells were incubated in [^13^C_5_]-Ino with or without Foro for 24 h, extracted, and analyzed as described in methods for PPP metabolites (B) and glycolytic/Krebs cycle metabolites (C) by IC-UHR-FTMS and by 1D HSQC NMR for inosine. All symbols and abbreviations are as described in **Figure 2**. Numbers in the X-axis represent those of ^13^C atoms in given metabolites. Values represent mean ± SEM (n=3). *, **, ***, and ***** denote p values ≤0.05, 0.01, 0.001, and 0.00001, respectively.

### Transformed cells display a diverse capacity for utilizing inosine as a carbon source

The nucleoside transporters and PNP are expressed in the majority of tissues and cell lines (data from The Human Protein Atlas). We therefore reasoned that cancer cells may also be able to take up and utilize inosine as an alternative carbon resource. To this end, we selected a panel of transformed cell lines and compared their growth rate in glucose-versus inosine-containing media. While most of the tested cell lines displayed some degree of inosine-dependent proliferation, several cell lines were unable to grow in inosine-containing media (Figure 5A, 5B and S6A). Next, we chose HeLa as a model cell line to study inosine catabolism and the role of PNP in inosine-dependent growth. We applied the same metabolic approach (Figure 2A) that was used to study T cells, to compare glucose and inosine catabolism in HeLa cells. HeLa cells displayed a similar pattern of ^13^C incorporation into intermediate metabolites in the central carbon metabolic pathways as in the T cells. Collectively, HeLa cells extensively catabolized [^13^C_6_]-Glc) and [^13^C_5_]-Ino, which derived fully ^13^C labeled isotopologues of intermediate metabolites in the PPP and glycolysis, and also a significant fraction of [^13^C_2_]-isotopologues in the Krebs cycle metabolites (Figure 5C and 5D). Consistent with the indispensable role of PNP in inosine catabolism in T cells, the pharmacological inhibition of PNP via Foro or genetic knockdown of PNP via siRNA significantly dampened inosine-dependent bioenergetics activity and cell growth in HeLa cells (Figure S6B and S6C). Together, these data suggest that some cancer cells are capable of utilizing inosine as an alternative fuel source to glucose in the central carbon metabolism.

**Figure 5.**
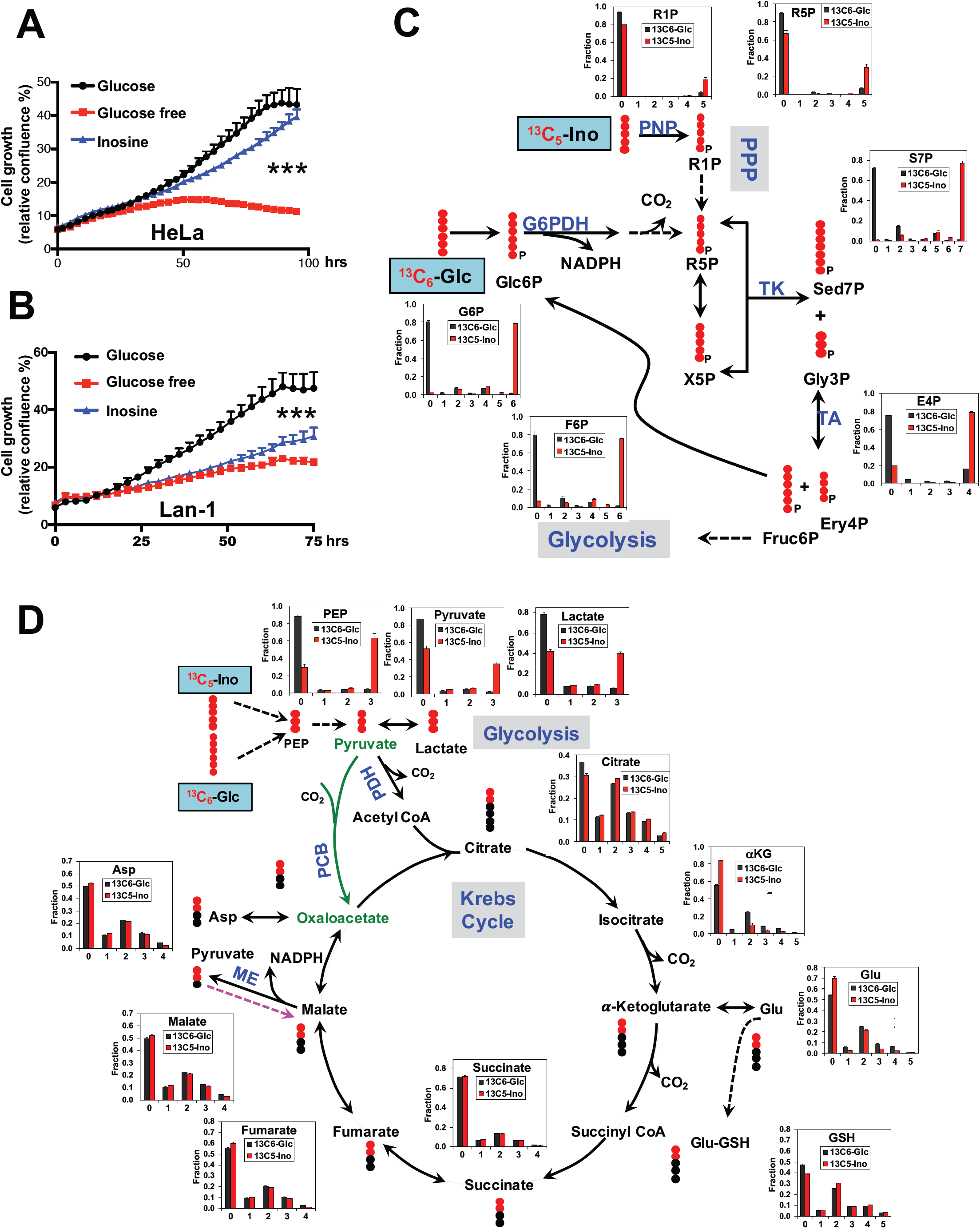
Transformed cells display diverse capacity to utilize inosine as a carbon source. (A) HeLa cell growth curves in the indicated conditional media were determined by live cell imaging analysis (IncuCyte ZOOM™). Error bars represent mean ± SD (n=4). ***, *P*<0.001 for 2 mM Inosine versus Glucose free. Data are representative of three independent experiments. (B) LAN-1 cells were cultured in the indicated conditional media and cell growth curves were monitored and analyzed by IncuCyte ZOOM™. Error bars represent mean ± SD (n=4). ***, *P*<0.001 for 2 mM Glucose versus 2 mM Inosine. Data are representative of three independent experiments. (C-D) HeLa cells were incubated with [^13^C_6_]-Glc or [^13^C_5_]-Ino for 24 h, extracted as described in Methods, and analyzed for PPP metabolites (C) and glycolytic/Krebs cycle metabolites (D) by IC-UHR-FTMS, respectively. All symbols and abbreviations are as described in **Figure 2**. Numbers in the X-axis represent those of ^13^C atoms in given metabolites. Values represent mean ± SEM (n=3).

### Inosine enhances the efficacy of T cell based immunotherapy in solid tumors

T_eff_ cells represent a key component of anti-tumor immunity. The checkpoint blockade approach that uses monoclonal antibodies targeting PD-1/PDL1 and CTLA4, and adoptive cell transfer (ACT) of tumor infiltrating lymphocytes (TILs) or CAR T cells are two front-line T cell-based immunotherapies [30; 42; 43; 44; 45; 46; 47; 48; 49]. Based on the *in vitro* findings described above, we reasoned that inosine supplement may improve T_eff_ cell-mediated anti-tumor activity, particularly toward tumor cells such as B16F10 or Lan-1 that are unable to utilize inosine to support cell growth (Figure S6A). PDL1 and PD-1 (PD) pathway blockade has been shown to elicit durable T cell-dependent anti-tumor responses. We thus assessed the anti-tumor effect of combining inosine and anti-PDL1 antibody in B16 melanoma bearing mice. While tumor growth, tumor-infiltrating T cells, and animal survival were comparable between vehicle (IgG control) and inosine treatment, the monotherapy of anti-PDL1 antibody clearly delayed tumor growth, prolonged animal survival time, and increased infiltrating T cells (Figure 6A and S7A). Importantly, mice treated with the combination of inosine and anti-PDL1 antibody displayed a better outcome (Figure 6A) and more tumor-infiltrating T cells than monotherapy group (Figure S7A). We also examined the ability of inosine to potentiate adoptive transfer based therapy which transfers Pmel+ T cells to mice bearing B16F10 xenografts (Figure 6B) or GD2-CAR T cells to immune-deficient mice bearing GD2 positive human neuroblastoma (LAN-1) xenografts (Figure 6C). While adoptively transferred T cells alone significantly delayed tumor growth and prolonged animal survival time, inosine plus T cells further potentiated these effects in the mouse xenograft models (Figure 6B, 6C and S7B). Taken together, our studies suggest that inosine supplementation enhances the potency and durability of T cell-based cancer immunotherapy in mouse preclinical models.

**Figure 6.**
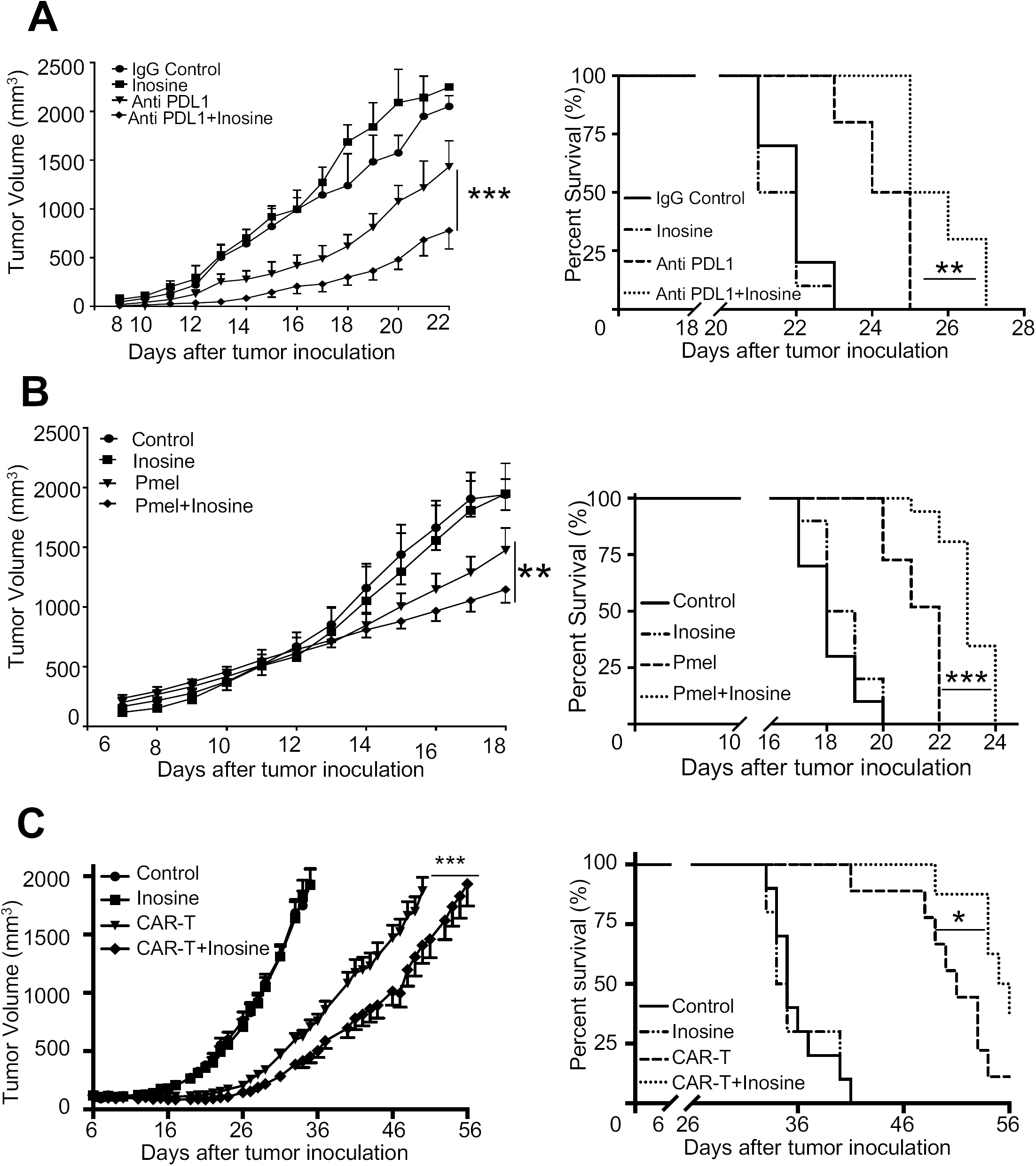
Inosine supplement enhances immunotherapy in targeting solid tumor. (A) Murine melanoma xenograft model was established in C57BL/6 mice by subcutaneously inoculation of B16F10 tumor cells. The indicated experimental mice were treated with IgG control (i.p at day 1, 4 and 7), inosine (oral gavage every day), anti-PDL1 antibody (i.p. at day 1, 4 and 7), and anti-PDL1 antibody (i.p. at day 1, 4 and 7) + inosine (oral gavage daily). Tumor size and mice survival were monitored. Error bars represent mean ± SEM (n=10). ***, *P*<0.001 for anti-PDL1 versus anti-PDL1+Inosine (left). **, *P*=0.0018 for anti-PDL1 versus anti-PDL1+Inosine (right). Data are representative of two independent experiments. (B) C57BL/6 mice were injected (s.c.) with B16F10 melanoma cells and sublethally irradiated (500cGy) at day 6 after tumor cell inoculation. One day later, mice were i.v. injected with activated Pmel CD8+ T cells (4×10^6^ cells/mouse). Mice were administered with Inosine 300 mg/kg/day by oral gavage from day 8 to 20. Tumor size and mice survival were monitored. Error bars represent mean ± SEM (n=10). **, *P*=0.0091 for Pmel versus Pmel+Inosine (left). ***, *P*<0.001 for Pmel versus Pmel+Inosine (right). Data are representative of two independent experiments. (C) Human neuroblastoma xenograft model was established in NSG mice by subcutaneously inoculation of LAN-1 neuroblastoma cells. The indicated experimental mice were treated with control (PBS i.v. at day 6 when tumors reach 100-150 mm^3^), inosine (300 mg/kg oral gavage daily starting from day 6), GD2-CAR-T cells (8×10^6^ cells/mouse i.v at day 6 when tumors grow to 100-150 mm^3^), and GD2-CAR-T cells (8×10^6^ cells/mouse i.v at day 6 when tumors grow to 100-150 mm^3^) + inosine (300 mg/kg oral gavage daily starting from day 6), respectively. Tumor growth and mice survival were monitored. Error bars represent mean ± SEM (n=16). ***, *P*<0.001 for CAR-T versus CAR-T+Inosine (left). *, *P*<0.05 for CAR-T versus CAR-T+Inosine (right). Data are representative of two independent experiments.

## Discussion

In addition to intrinsic cell signaling cascade, numerous extrinsic environmental factors including oxygen and tissue-specific nutrient supplies significantly influence T cell metabolic programming and thus activation. T cells are distributed throughout the body and thus encounter variable metabolic stresses depending on where tumors or infections occur. A key factor could be the competition for nutrients between infiltrating T cells and rapidly proliferating cells including cancer cells or pathogens. It is likely that a wide range of abundant bioenergetic carbohydrates in blood plasma could be a source of nutrients for proliferative T cells and cancer cells *in vivo* [91; 92; 93]. Mounting evidence suggests that cancer cells can also scavenge extracellular protein or utilize acetate and ketone bodies as alternative fuels [94; 95; 96]. Although glucose is considered as the primary fuel for proliferating T cells, galactose can replace glucose to support T cell survival and proliferation following activation [97]. Our studies suggest that T cells, to various extents, can metabolize other naturally occurring saccharides or their derivatives, such as D-fructose and D-mannose, as fuels for supporting survival and proliferation. Our studies also revealed T cell’s capacity to utilize inosine for energy and effector cell functions. Such metabolic plasticity is likely to be crucial to T cell’s ability to elicit robust immune responses in different tissue contexts.

Although D-ribose produced via the ribose salvage pathway from nucleotide or nucleoside can serve as a carbon and bioenergetic source under nutrient starvation, proliferative normal or transformed cells are unable to catabolize free ribose due to the lack of functional ribose transporters or key enzymes [98; 99; 100; 101; 102]. In contrast, the expression of phosphorylase (PNP) that can convert inosine into ribose-1-phosphate and hypoxanthine is consistent with our finding on inosine’s capacity as a ribose donor in T cells. Such PNP-mediated breakdown of inosine is energetically more efficient than free ribose metabolism, as it only needs inorganic phosphate while the latter requires ATP. We found that the ribose derived from inosine was catabolized efficiently through the central carbon metabolic routes including the PPP, glycolysis, and the Krebs cycle in place of glucose. This, together with inosine’s ability to support T_eff_ cell growth and functions in the absence of glucose, suggests that inosine, as a ribose donor, may replace glucose as a key fuel source for T_eff_ cells under glucose deprivation, as is found in solid tumors.

The degradation of nucleotides and nucleosides is an evolutionarily conserved metabolic process, which can supply carbon and energy to support cell proliferation and functions [98; 99; 100; 101; 102; 103; 104; 105; 106; 107]. However, we found that inosine but no other nucleosides (adenosine and guanosine) supported T cell proliferation. This result suggests that inosine is a preferred substrate of nucleoside transporters and nucleoside phosphorylase that are predominantly expressed in T cells. Furthermore, PNP inhibitor abolished inosine uptake and metabolism as well as its ability to support T_eff_ cell proliferation and functions *in vitro*. Several previous studies also demonstrated the role of inosine in supporting bioenergetic and function in human red blood cells, swine and chicken erythrocytes, which lack glucose transporters, as well as enhancing the recovery of ischemic heart and kidney by preserving ATP [108; 109; 110; 111; 112; 113; 114; 115].

Inosine and adenosine can maintain ATP during hypoxia and nutrient restriction in the central nerve system (CNS) and the catabolism of the ribose moiety of inosine is necessary to provide the bioenergetic support for CNS cells [116]. The protective effect of adenosine depends on adenosine deaminase (ADA), which converts adenosine into inosine [117]. Although inosine and adenosine concentrations in blood plasma is in the low micromolar range, they can accumulate to hundreds of micromolar via catabolism of nucleotides and nucleosides in the necrotic, hypoxic, or inflammatory microenvironment, which is characteristic of solid tumors [118; 119; 120; 121; 122; 123]. T cells may utilize inosine and adenosine accumulated in the tumor microenvironment (TME) as an important carbon and energy source.

In response to tissue injury, stressed or damaged cells release ATP and its product, adenosine, into the extracellular space to elicit autocrine and paracrine signaling responses through cell-surface P2 purinergic receptors [85; 124; 125]. To terminate adenosine-mediated signaling, adenosine deaminase (ADA) converts adenosine to inosine, rendering inosine a weak agonist of adenosine receptor [126]. Consistent with these findings, we found that inosine but not adenosine enhances T_eff_ cell functions, which may be mediated via inosine catabolism, as the inhibitor of PNP abolished both inosine catabolism and inosine-induced activation of T_eff_ cell functions. However, it is still conceivable that some of the *in vivo* effects of inosine are mediated by purinergic signaling due to the concurrent depletion of immune suppressive adenosine. We therefore surmise that ADA treatment may reduce immune suppressive adenosine in TME while increasing the levels of inosine to fuel tumor infiltrated T cells, thereby improving anti-tumor immunity. The genetic disorder leading to the loss of ADA enzymatic function can cause severe combined immunodeficiency (SCID) [127; 128]. As such, the polyethylene glycol–conjugated adenosine deaminase (PEG-ADA) and ADA gene therapy have been developed as the first enzyme replacement therapy (ERT) to treat ADA-SCID [129; 130]. Based on our findings, further research on the molecular action of PEG-ADA and its repurposing in cancer immunotherapy is warranted.

Anti-tumor immunotherapy not only offers the potential of high selectivity in targeting and killing tumor cells, but also represents an appealing addition to the current therapeutic regimens due to its potential to eradicate recurrent/metastatic disease following conventional therapies. However, the immunosuppressive microenvironment is a major barrier to the development of effective immunotherapy. It has been shown that metabolic modulation of T cells is a valid approach to alter T cell mediated immune responses in a variety of physio-pathological contexts [15; 16; 24; 67; 68; 69; 75; 131; 132; 133; 134; 135; 136; 137]. Clearly, nutrient restriction in solid tumor can play an important role in restraining anti-tumor activity of infiltrating T cells. We have shown that inosine readily replaced glucose in supporting T_eff_ cells growth and functions *in vitro*, and inosine supplement improved T_eff_ cell-mediated anti-tumor activities in animal models. Inosine supplementation via oral administration or intravenous infusion has been shown to be safe and tolerable in recent clinical trials of inosine supplementation in Multiple Sclerosis and Parkinson Disease [138; 139; 140; 141]. We therefore envision that metabolic modulation of T cells including inosine and PEG-ADA supplementation poses a new complementary strategy to optimize the potency and durability of cancer immunotherapy. Additional studies on the formulation of inosine supply or strategies that may enhance the local accumulation of inosine in the TME are warranted to facilitate future clinical development of metabolic modulation in cancer immunotherapy.

## ACKNOWLEDGEMENTS

This work was supported by 1R21CA227926-01A1, 1UO1CA232488-01 and 1R01AI114581 from National Institute of Health, V2014-001 from the V-Foundation and 128436-RSG-15-180-01-LIB from the American Cancer Society (to RW); 1P01CA163223-01A1, 1U24DK097215-01A1 (to TWMF, ANL); Redox Metabolism Shared Resource(s) of the University of Kentucky Markey Cancer Center (P30CA177558). ^13^C-enriched standards were obtained from NIH Common Fund Metabolite Standards Synthesis Core (http://www.metabolomicsworkbench.org/standards/index.php). We thank John Sherman for critically reading and editing the manuscript.

## Material and Methods

### Mice

C57BL/6 and NSG mice were purchased from Envigo (formly Harlan) and the Jackson Laboratory. Pmel transgenic mice (B6.Cg-Thy1^a^/Cy Tg(TcraTcrb)8Rest) were purchased from the Jackson Laboratory [79]. Mice at 7–12 weeks of age were used. All mice were kept in specific pathogen – free conditions within the Animal Resource Center at the Research Institute at Nationwide Children’s Hospital and Baylor College of Medicine. Animal protocols were approved by the Institutional Animal Care and Use Committee of the Research Institute at Nationwide Children’s Hospital and Baylor College of Medicine.

### Tumor cell culture and reagents

LAN-1, a GD2-positive tumor cell line, and B16-F10 cell lines were purchased from ATCC and were grown in RPMI 1640 and DMEM media (Corning Cellgro, Thermo Fisher Scientific, Grand Island, NY) with 10% fetal calf serum (Gibco-Invitrogen, Carlsbad, CA) and 1% penicillinstreptomycin (Corning) in a 37 °C humidified atmosphere of 95% air and 5% CO_2_, respectively. HeLa (human cervical cancer cells, ATCC) cells were grown at 37 °C/5% CO_2_ in DMEM media supplemented with 10% fetal calf serum and 1% penicillin-streptomycin. Glucose-free DMEM was supplemented with 10% (v/v) heat-inactivated dialyzed fetal bovine serum, which was made dialyzing against 100 volumes of distilled water (five changes in 3 days) using Slide-ALyzer™ G_2_ dialysis cassettes with cut-through MW size 2K (Thermo Fisher Scientific) at 4 °C. Forodesine Hydrocholoride was purchased from MedChem Express (Monmouth Junction, NJ). Anti-PNP and anti-actin antibodies were from Santa Cruz Biotechnology (Santa Cruz, CA, USA).

### Human and mouse T cells isolation and culture

PBMCs were collected from healthy donors, approved by Baylor College of Medicine review board. Mononuclear cells from peripheral blood (PBMCs) were isolated by Ficoll/Hypaque density gradient centrifugation. In some experiment, human T cells were directly enriched from PBMC by negative selection using MojoSort™ Human CD3 T Cell Isolation Kit (Biolegend, CA) following the manufacturer’s instructions. Isolated human mononuclear cells or enriched human T cells were either maintained in culture media containing 10 ng/ml IL-7 or stimulated human IL-2 (100 U/ml) and plate-bound anti-CD3 and anti CD28 antibody. Plates were pre-coated with 1 μg/ml antibodies overnight at 4 °C. To generate GD2-CAR T cells, activated human T cells were transduced with GD2-CAR retrovirus on retronectin-coated nontissue culture plates and cultured with IL-2 for one week [81]. Mouse total T and CD8+ native T cells were enriched from spleens and lymph nodes by negative selection using MACS system (MiltenyiBiotec, Auburn, CA) following the manufacturer’s instructions. For total T cells culture, freshly isolated total T cells with 75-80% CD3 positivity were either maintained in culture media containing 5 ng/ml IL-7 or were stimulated with human IL-2 (100 U/ml) as well as plate-bound anti-mCD3 (clone 145-2C11) and anti-mCD28 (clone 37.51) antibodies. Plates were pre-coated with 2 μg/mL antibodies overnight at 4 °C. For mouse and human T cell CFSE dilution analysis, 10-20 x10^6^ cells were preincubated for 10 min in 4 μM CFSE (Invitrogen) diluted in PBS plus 5% FBS before culture. For CD8+ T cells activation and culture, naive CD8 T cells were activated by plate-bound antibodies and incubated with recombinant mouse IL-2 (100 U/mL) and IL-12 5 ng/mL for 5 days. For Pmel+ CD8+ T cell activation and culture, splenocytes from Pmel were isolated and culture with 1µM human gp100 (hgp100) (Genscript) and 30 IU/mL recombinant human IL-2 (PeproTech Inc) in complete media for 5-7 days [142]. The cells were then cultured in RPMI 1640 media supplemented with 10% (v/v) heat-inactivated fetal bovine serum (FBS), 2 mM L-glutamine, 0.05 mM *2-mercaptoethanol*, 100 units/mL penicillin and 100 μg/mL streptomycin at 37°C in 5% CO_2_. Glucose free RPMI 1640 media that was supplemented with 10% (v/v) heat-inactivated dialyzed fetal bovine serum (DFBS) was used as basal conditional media for glucose and inosine reconstitution. DFBS was made dialyzing against 100 volumes of distilled water (five changes in three days) using Slide-ALyzer™ G2 dialysis cassettes with cut-through MW size 2K (ThermoFisher Scientific) at 4°C.

### Flow cytometry

For the analysis of surface markers, cells were stained in PBS containing 2% (w/v) BSA and the appropriate antibodies listed in table S1. For intracellular cytokine staining, T cells were stimulated for 4-5 h with phorbol 12-myristate 13-acetate (PMA) and ionomycin in the presence of monensin before being stained according to the manufacturer’s instructions (BD Bioscience). Flow cytometry data were acquired on Novocyte (ACEA Biosciences) or LSRII (Becton Dickinson) and were analyzed with FlowJo software (TreeStar). All the chemicals and antibodies used are listed in Table S1.

### Metabolite screening and bioenergetics analysis

Metabolite screening was performed using 96-well PM-M1 and PM-M2 phenotyping microarray for mammalian cells (BioLog Inc). Human or mouse activated T cells were washed twice with PBS, and resuspended in glucose and phenol red free DMEM media supplemented with 10% dialyzed FBS and 100 U/mL IL-2 followed by immediate inoculation of 100 *µ*L cells/well (1×10^6^ cells/ml) into the Biolog plates. After 24 h incubation at 37°C in a 5% CO_2_ incubator, 20 *µ*L of Redox Dye Mix MB (Biolog Inc) was added to each well, and absorbance at 590 and 750 nm wavelengths were recorded at different time points.

### CD8+ CTL tumor killing assay

CTL tumor killing assay was performed using caspase-3/7 reagent (Essen BioScience, Ann Arbor, MI) and the IncuCyte ZOOM live-cell imaging system (Essen BioScience) which is placed in an incubator at 37 °C and 5% CO_2_. LAN-1 cells, (1.5×10^4^ cells /well) and B16F10 (1.5×10^3^ cells/well) were plated in 96 well plates and allowed to grow until 20% confluence was achieved. The media was replaced and target cells were co-cultured with GD2-CAR T cells or hgp100 activated Pmel T cells respectively in 10:1 (effector: target) ratios in glucose free media or with 2 mM glucose or Inosine or adenosine, and caspase 3/7 reagent was added at 1:1000 dilution to each well. Dead cells, which are positive for Caspase-3/7, were monitored and quantified automatically acquired at regular intervals by the IncuCyte software.

### Tumor xenograft and T cell *in vivo* imaging

For the LAN-1 xenograft, 1.5×10^6^ LAN-1 tumor cells were mixed in 100 μl matrigel (Corning) and were subcutaneously inoculated in the dorsal left and right flanks of NSG mice. 8 x10^6^ GD2-CAR-T or GD2-CAR-T-Luciferase cells were intravenously injected into tumor-bearing mice when tumor grew to about 4-6 mm in diameter (around 6-8 days). For inosine treatments, inosine (Sigma) was administered (300 mg/kg in PBS) by oral gavage daily after CAR-T cell administration and throughout the experiment. For T cell *in vivo* imaging, the images were captured using IVIS imaging system (Xenogen) after i.v injection of 150 mg/kg D-luciferin (Xenogen, Alameda, CA) at day 4 and day 7 after GD2-CAR-T-Luciferase cells administration. Photon emission was analyzed by constant region-of-interest (ROI) drawn over the tumor region and the signal measured as total photon/sec/cm2/steridian (p/s/cm2/sr).

For B16F10 melanoma model, female C57BL6 mice were inoculated with 1×10^5^ cells in the flank subcutaneously at day 0 and treated on days 1 (200 μg), 4 and 7 (100 μg) with anti-PDL1 antibody. Inosine (300 mg/kg) was administered by oral gavage every day, until animals reach end point and tumor volume (mm^3^) and overall survival were assessed throughout the experiment. To evaluate the tumor infiltrating immune cells, after 10 and 15 days of inosine treatment, tumors were dissected and dissociated using gentle MACS™ Dissociators according to manufacturer’s instructions. Cells were stained with surface antibodies and analyzed using flow cytometry. In adoptive transfer experiments, C57BL/6 mice were injected (s.c.) with 1X10^5^ B16F10 melanoma cells and animals were irradiated sublethaly (500cGy) after 6 days of tumor cell inoculation. On day 7, 4X10^6^ active Pmel T cells were injected (i.v.) and inosine (300 mg/kg oral gavage) was administered daily and tumor size was monitored until animals reach the endpoint.

### Metabolite extraction and analysis by ion chromatography-ultra high resolution-Fourier transform mass spectrometry (IC-UHR-FTMS) and NMR

Cells were cultured in glucose-free media with 2 mM ^13^C_6_-glucose (Cambridge Isotope Laboratories) or [^13^C_5_]-ribose-inosine (Omicron Biochemicals) for 24 h at 37 °C and were then washed 3 times in cold PBS before metabolic quenching in cold acetonitrile and harvested for metabolite extraction as described previously [143]. The polar extracts were reconstituted in nanopure water before analysis on a Dionex ICS-5000+ ion chromatography system interfaced with a Thermo Fusion Orbitrap Tribrid mass spectrometer (Thermo Fisher Scientific) as previously described [144] using a *m/z* scan range of 80-700. Peak areas were integrated and exported to Excel via the Thermo TraceFinder (version 3.3) software package before natural abundance correction [145]. The isotopologue distributions of metabolites were calculated as the mole fractions as previously described [146]. The number of moles of each metabolite was determined by calibrating the natural abundance-corrected signal against that of authentic external standards. The amount was normalized to the amount of extracted protein, and is reported in nmol/mg protein.

Polar extracts reconstituted in D_2_O (> 99.9%, Cambridge Isotope Laboratories, MA) containing 0.5 mmole/L d_6_-2,2-dimethyl-2-silapentane-5-sulfonate (DSS) as internal standard were analyzed by 1D ^1^H and ^1^H{^13^C}-HSQC NMR on a 14.1 T DD2 NMR spectrometer (Agilent Technologies, CA). 1D ^1^H spectra were acquired using the standard PRESAT pulse sequence with 512 transients, 16384 data points, 12 ppm spectral width, an acquisition time of 2 s and a 6 s recycle time with weak irradiation on the residual HOD signal during the relaxation delay. The raw fids were zero filled to 131072 points and apodized with 1 Hz exponential line broadening prior to fourier transformation. 1D HSQC spectra were recorded with an acquisition time of 0.25 s with GARP decoupling, and recycle time of 2 s over a spectral width of 12 ppm, with, 1024 transients. The HSQC spectra were then apodized with unshifted Gaussian function and 4 Hz exponential line broadening and zero filled to 16k data points before Fourier transformation. Metabolites were assigned by comparison with in-house [147] and public NMR databases. Metabolite and their ^13^C isotopomers were quantified using the MesReNova software (Mestrelab, Santiago de Compostela, Spain) by peak deconvolution. The peak intensities of metabolites obtained were converted into nmoles by calibration against the peak intensity of DSS (27.5 nmoles) at 0 ppm for ^1^H spectra and that of phosphocholine at 3.21 ppm (nmoles determined from 1D ^1^H spectra) for HSQC spectra before normalization with mg protein in each sample.

### siRNA transfection in HeLa cells

The siRNA oligonucleotides corresponding to human PNP (Purine nucleoside phosphorylase) were purchased from Fisher. siRNA oligonucleotides (20 nM) were transfected into HeLa cells using Lipofectamine RNAiMAX reagent (Invitrogen). After 72 h of transfection, immunoblots were carried out to examine the knockdown of targeted proteins.

### Electroporation of human T cells

PBMC cells were stimulated with plate-bound anti-CD3 and anti-CD28 antibodies for two days before electroporation. Cells were centrifuged and washed once with PBS. 1×10^6^ cells were resuspended in 100 μL electroporation buffer T (Thermo Fisher, Waltham, MA) and 100 nM siRNA oligonucleotides corresponding to human PNP was added. Electroporation was performed at 2150 V 20 ms 1 pulses settings for stimulated PBMC cells using Neon electroporation device (Thermo Fisher, Waltham, MA). Immediately after electroporation, the cells were plated in 12-well plate with 100 IU/ml IL-2 and incubated at 37°C.

### qPCR and western blot analysis

Total RNA was isolated using the Quick-RNA™ MiniPrep Kit (Zymo Research) and was reverse transcribed using random hexamer and M-MLV Reverse Transcriptase (Invitrogen). SYBR green-based quantitative RT-PCR was performed using the BIO-RAD CFX96™Real-Time System. The relative gene expression was determined by the comparative *C*_T_ method also referred to as the 2^-ΔΔ*C*^ _T_ method. The data were presented as the fold change in gene expression normalized to an internal reference gene (beta2-microglobulin) and relative to the control (the first sample in the group). Fold change=2^-ΔΔ*C*^_T_=[(*C*_Tgene of interst_ - *C*_Tinternal reference_)]sample A-=[(*C* _Tgene of interst_ - *C* _Tinternal reference_)]sample B. Samples for each experimental condition were run in triplicate PCR reactions. Primer sequences were obtained from PrimerBank [148]. The following pair of primers were used for detecting mouse PNP (5’-ATCTGTGGTTCCGGCTTAGGA-3’ and 5’-TGGGGAAAGTTGGGTATCTCAT-3’). Cell extracts were prepared and immunoblotted as previously described [149].

### Metabolic activity analysis

Fatty acid β-oxidation flux was determined by measuring the detritiation of [9,10^−3^H]-palmitic acid [150; 151]. One million T cells were suspended in 0.5ml fresh media. The experiment was initiated by adding 3 *µ*Ci [9,10^−3^H]-palmitic acid complexed to 5% BSA (lipids free; Sigma) and, 2 h later, media were transferred to 1.5 mL microcentrifuge tubes containing 50 μL 5 N HCL. The microcentrifuge tubes were then placed in 20 mL scintillation vials containing 0.5 mL water with the vials capped and sealed. ^3^H_2_O was separated from unmetabolized [9,10^−3^H]-palmitic acid by evaporation diffusion for 24 h at room temperature. A cell-free sample containing 3 *µ*Ci [9,10^−3^H]-palmitic acid was included as a background control.

Gluamine oxidation activity was determined by the rate of ^14^CO_2_ released from [U-^14^C]-glutamine [152]. In brief, one-five million T cells were suspended in 0.5 ml fresh media. To facilitate the collection of ^14^CO_2_, cells were dispensed into 7ml glass vials (TS-13028, Thermo) with a PCR tube containing 50μl 0.2M KOH glued on the sidewall. After adding 0.5 μci [U-^14^C]-glutamine, the vials were capped using a screw cap with rubber septum (TS-12713, Thermo). The assay was stopped 2h later by injection of 100μl 5N HCL and the vials were kept at room temperate overnight to trap the ^14^CO_2_. The 50μl KOH in the PCR tube was then transferred to scintillation vials containing 10ml scintillation solution for counting. A cell-free sample containing 0.5μci [U-^14^C]-glutamine was included as a background control.

#### Statistical analysis

*P* values were calculated with Student’s *t*-test. *P* values smaller than 0.05 were considered significant, with *P*-values<0.05, *P*-values<0.01, and *P*-values<0.001 indicated as *, **, and ***, respectively.

## Figure legends

**Figure S1.**
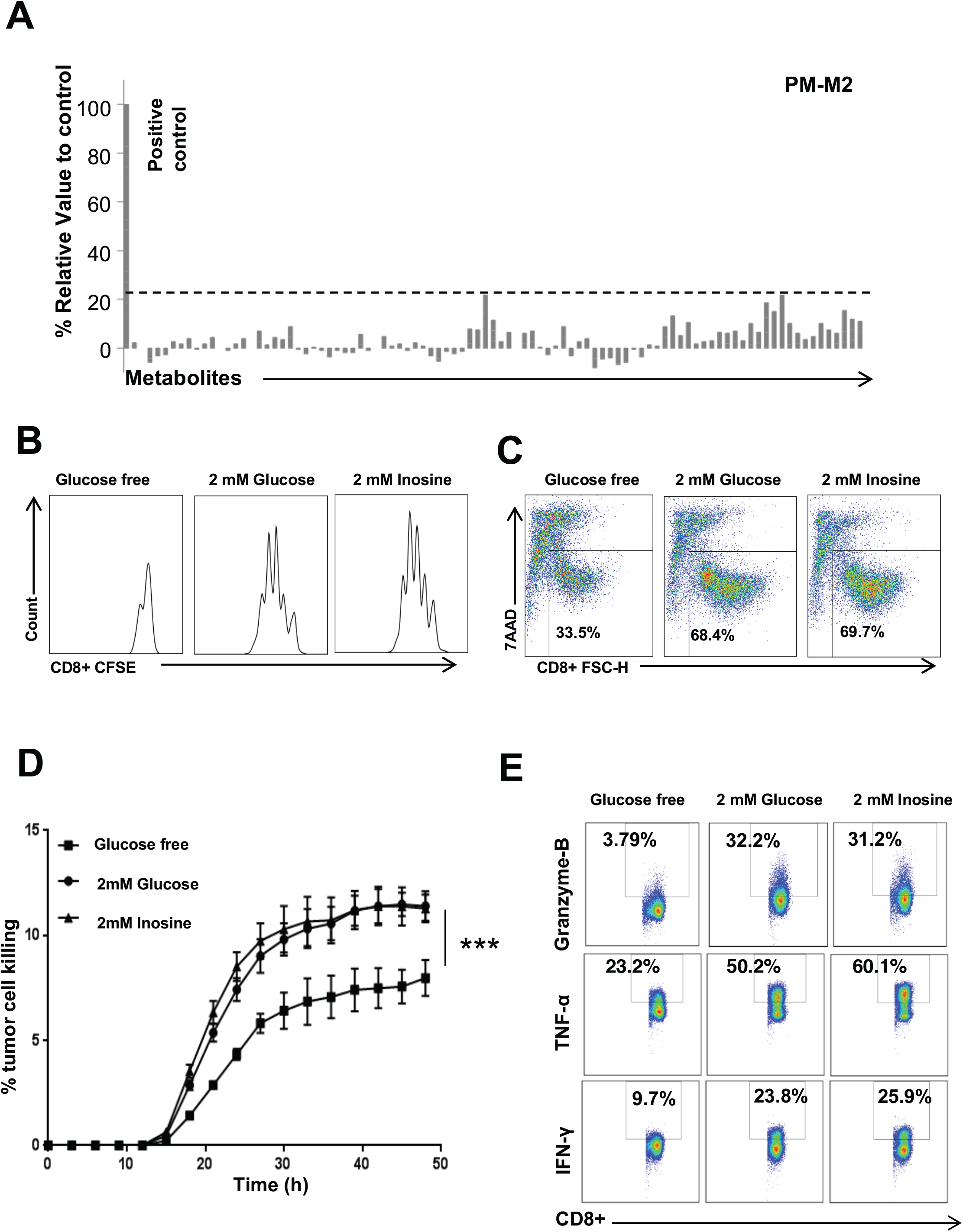
Inosine can support T effector cells proliferation and function in the absence of glucose. (A) Metabolite array from the PM-M2 Microplate (Biolog). Activated human T cells were incubated with various energy sources in a PM-M2 Microplate for 24 h, followed by Biolog redox dye mix MB incubation and measured spectrophotometrically at 590 nm. (B-C) Naive CD8+ T cells from human PBMC were activated by plate-bound anti-CD3 and anti-CD28 antibodies in the indicated conditional media for 5 days. Cell proliferation and cell death of active CD8+ T cells were determined by CFSE dilution and 7AAD uptake, respectively. Data are representative of three independent experiments. (D) LAN-1 cells were co-cultured with GD2-CAR T cells in the indicated conditional media and cell death of LAN-1 was monitored using caspase 3/7 fluorescent substrate by live cell imaging and quantified using the IncuCyte ZOOM™ analyzer. Error bars represent standard deviation (n=4). ***, *P*<0.001 for glucose free versus 2 mM Inosine. (E) The indicated proteins in GD2-CAR T cells were quantified by intracellular staining following PMA and ionomycin stimulation.

**Figure S2.**
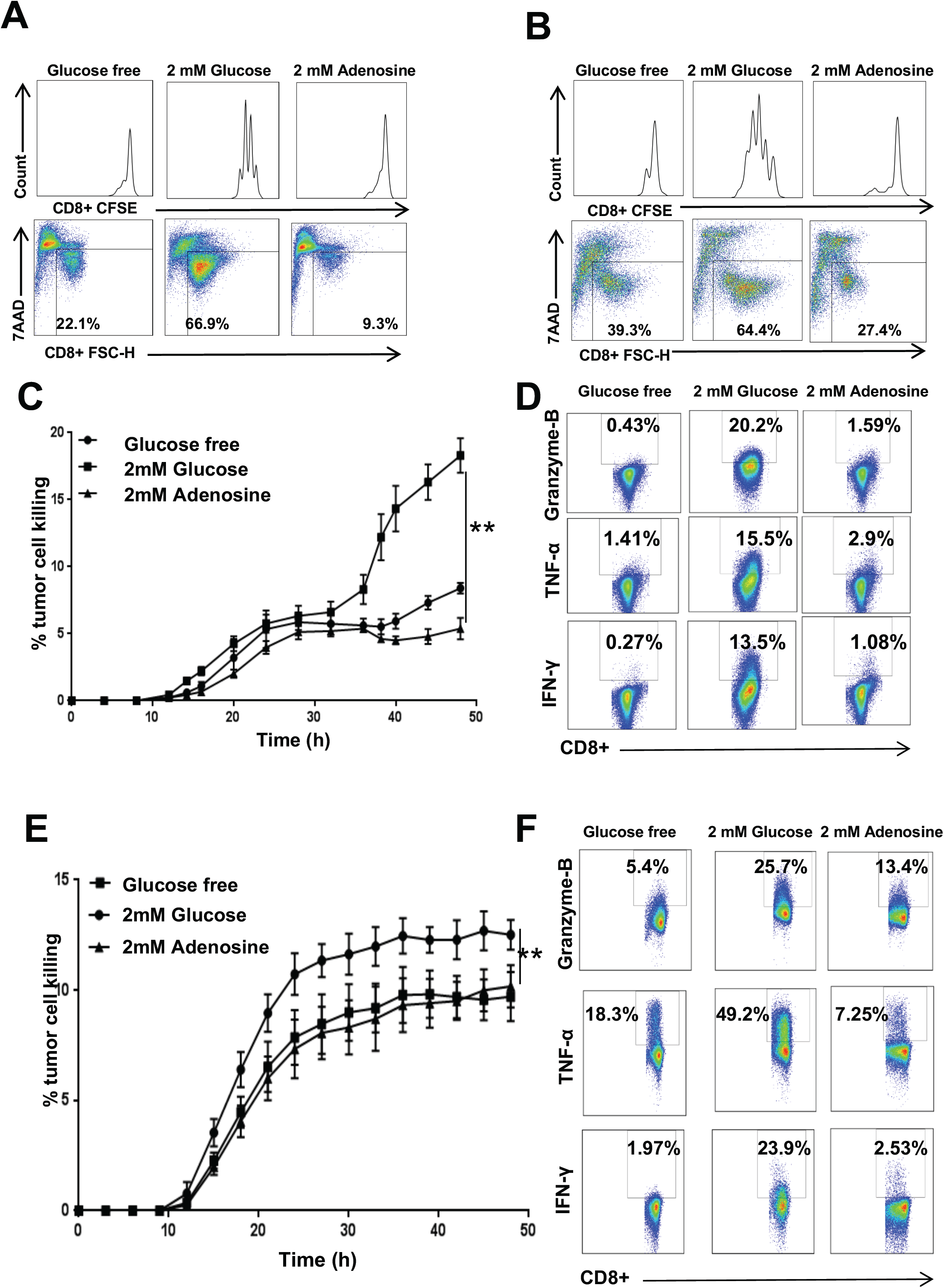
Adenosine cannot support T effector cells proliferation and function in the absence of glucose. (A-B) Naive CD8+ T cells from C57BL/6 mice or from human PBMC were activated by plate-bound anti-CD3 and anti-CD28 antibodies in the indicated conditional media for 3-5 days. Cell proliferation and cell death of active CD8+ T cells were determined by CFSE dilution and 7AAD uptake, respectively. Data are representative of three independent experiments. (C) B16F10 melanoma cells were co-cultured with activated Pmel T cell in the indicated conditional media, and tumor cell death was evaluated using caspase 3/7 reagent by IncuCyte ZOOM™. Error bars represent standard deviation (n=4). **, *P* =0.007 for 2 mM glucose versus 2 mM Adenosine. Data are representative of two independent experiments. (D) Naive CD8+ T cells from C57BL/6 mice were activated by plate bound anti-CD3 and anti-CD28 antibodies and differentiated in the indicated conditional media for 4 days. The indicated proteins were quantified by intracellular staining following PMA and ionomycin stimulation. (E) LAN-1 cells were co-cultured with GD2-CAR T cells in the indicated conditional media and cell death of LAN-1 was monitored using caspase 3/7 fluorescent substrate by IncuCyte ZOOM™. Error bars represent standard deviation (n=4). ***, *P*<0.001 for 2 mM Glucose versus 2 mM Adenosine. Data are representative of two independent experiments. (F) The indicated effector molecules in GD2-CAR T cells were quantified by intracellular staining following PMA and ionomycin stimulation.

**Figure S3.**
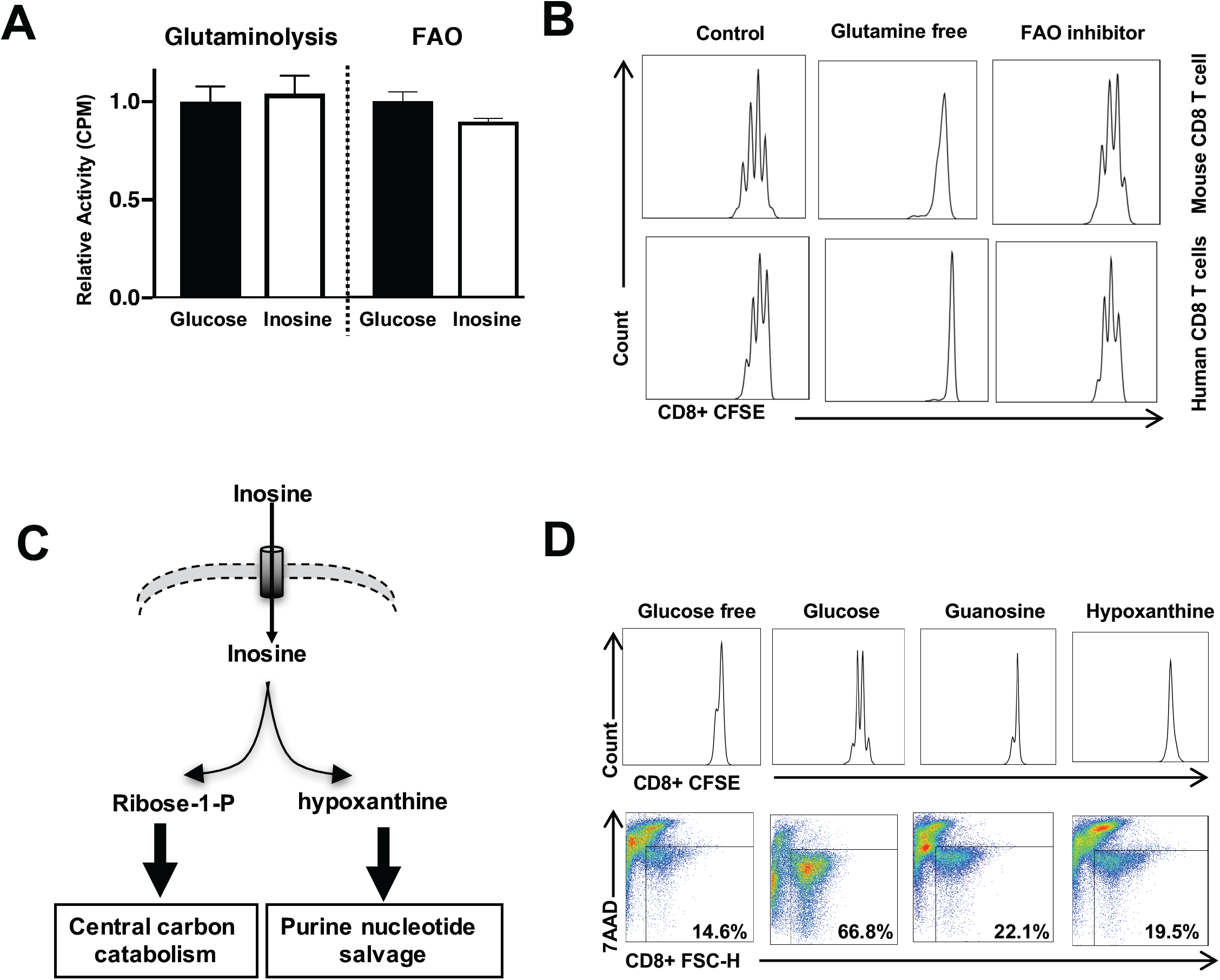
Inosine does not support T effector cell proliferation by enhancing glutamine and fatty acids catabolism or through purine nucleotide salvage. (A) Naive CD8+ T cells from C57BL/6 mice were activated by plate-bound anti-CD3 and anti-CD28 antibodies in complete media for 24 h, then the cells were switched to indicated conditional media and were cultured for 24 h. Metabolic fluxes of glutaminolysis and fatty acid β-oxidation (FAO) were measured by the generation of ^14^CO_2_ from [U-^14^C]-glutamine and ^3^H_2_O from from [9, 10^−3^H]-palmitic acid, respectively. Error bars represent standard deviation from the mean of triplicate samples. Data are representative of two independent experiments. (B) Naive CD8+ T cells from C57BL/6 mice (upper) and human PBMC (lower) were activated by plate-bound anti-CD3 and anti-CD28 antibodies in complete media for 24 h, then cells were switched to indicated culture condition for 3-5 days. Cell proliferation of active CD8+ T cells was determined by CFSE dilution. Data are representative of two independent experiments. (C) Diagram of inosine uptake and breakdown into ribose-1-phosphate (Ribose-1-P) and hypoxanthine, entering central carbon catabolism and the purine nucleotide salvage pathway, respectively. (D) Naive CD8+ T cells from C57BL/6 mice were activated by plate-bound anti-CD3 and anti-CD28 antibodies for 24 h, then cells were switched to indicated conditional media for 72 h. Cell proliferation and cell death were determined by CFSE dilution (upper) and 7AAD uptake (lower), respectively. Data are representative of three independent experiments.

**Figure S4.**
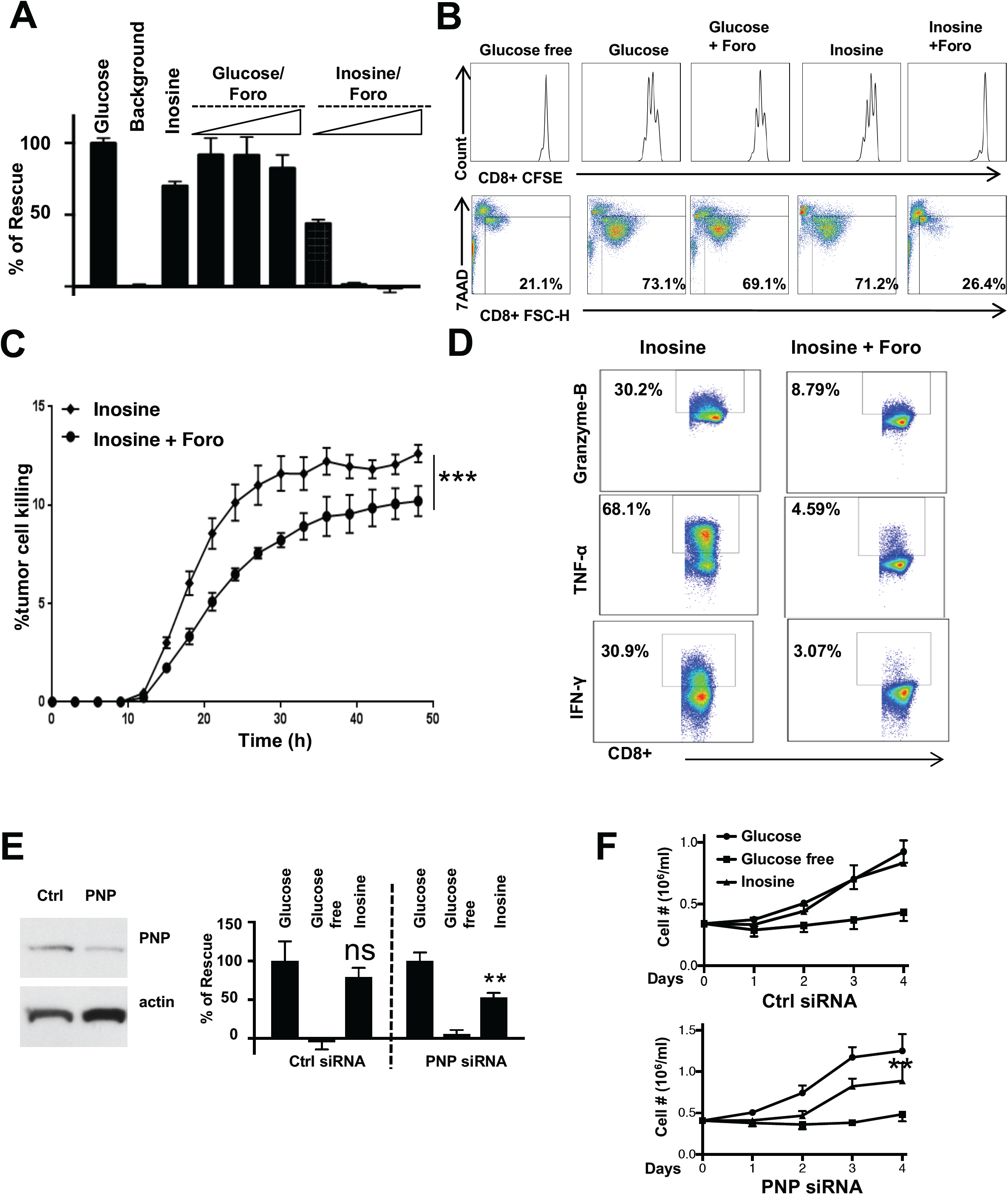
The purine nucleoside phosphorylase (PNP) is required for inosine-dependent human T effector cell proliferation and effector functions. (A) Activated human T cells were incubated without glucose (background), with glucose or with inosine in combination with increased concentrations of Foro (0.1 μM, 0.5 μM, and 2 μM) for 24 h, followed by Biolog redox dye mix MB incubation and measured spectrophotometrically at 590 nm. (B) Naive CD8+ T cells from human PBMC were activated by plate-bound anti-CD3 and anti-CD28 antibodies in the indicated conditional media with or without 2 μM Foro for 5 days. Cell proliferation and cell death of active CD8+ T cells were determined by CFSE dilution (upper) and 7AAD uptake (lower), respectively. Data are representative of three independent experiments. (C) LAN-1 cells were co-cultured with GD2-CAR T cells in the presence of inosine with or without Foro and cell death of LAN-1 was determined by using caspase 3/7 fluorescent substrate with live cell imaging analysis (IncuCyte ZOOM™). Error bars represent standard deviation (n=4). ***, *P*<0.001 for 2 mM Inosine versus 2 mM Inosine + 2 μM Foro. (D) The indicated proteins were quantified by intracellular staining following PMA and ionomycin stimulation. Data are representative of three independent experiments. (E) Human T cells from human PBMC were activated by plate-bound anti-CD3 and anti-CD28 antibodies for 2 days. Activated human T cells were electroporated with scrambled siRNA or PNP siRNA using Neon system and cultured for 2 days. PNP protein expression levels were determined by immunoblot (left). Cells were then incubated without glucose (background), with glucose or with inosine for 24 h, followed by Biolog redox dye mix MB incubation for 2 h and measured spectrophotometrically at 590 nm (right). (F) Proliferation curves of human T cells transfected with scrambled siRNA (upper) or PNP siRNA (lower) cultured in the indicated conditional media were determined by counting cell number daily. Error bars represent standard deviation (n=3). **, *P*<0.01 for 2 mM Glucose versus 2 mM Inosine in PNP siRNA transfected cells (lower). Data are representative of three independent experiments.

**Figure S5.**
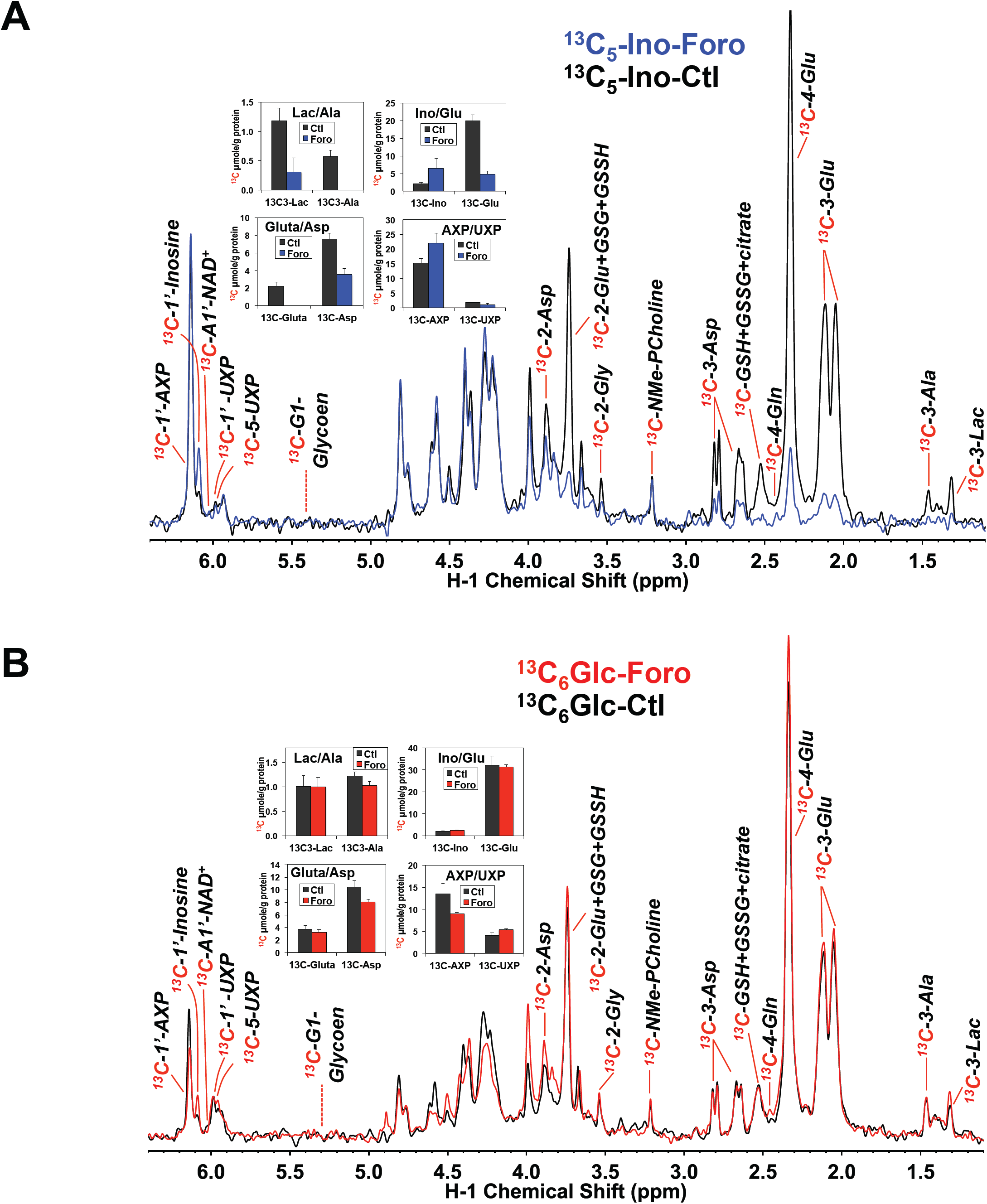
PNP inhibitor forodesine blocks ^13^C incorporation from [^13^C_5_]-ribose-inosine into central metabolites but has little effect on [^13^C_6_]-glucose catabolism. The same experiment as in **Figure 4** was performed for active human T cells ± Foro in the presence of [^13^C_5_]-inosine or [^13^C_6_]-Glc for 24 h. The polar extracts from both tracer experiments were analyzed by 1D HSQC NMR for ^13^C abundance (spectra as shown). Selected metabolites were quantified as ^13^C µmoles/g protein using the N-methyl resonance of phosphocholine (NMe-PCholine) as an internal standard, which are shown as bar graphs for the [^13^C_5_]-inosine (A) and [^13^C_6_]-glucose tracer (B) experiments. Values are average ± SEM (n=3). Dashed line denotes the expected peak position of the H1-glucose resonance of glycogen. The proton resonances used for quantification were: ^13^C_3_-Lac/Ala – the H3 resonances of lactate/Ala; ^13^C-Gluta – the H4-Glu resonance of GSH + GSSG with minor contribution of H2,5 resonance of citrate; ^13^C-Ino, ^13^C-Glu, ^13^C-Asp, ^13^C-AXP, and ^13^C-UXP – H1’, H4, H3, and H1’ resonances of inosine, Glutamate, Aspartate, and adenine/uracil nucleotides, respectively.

**Figure S6.**
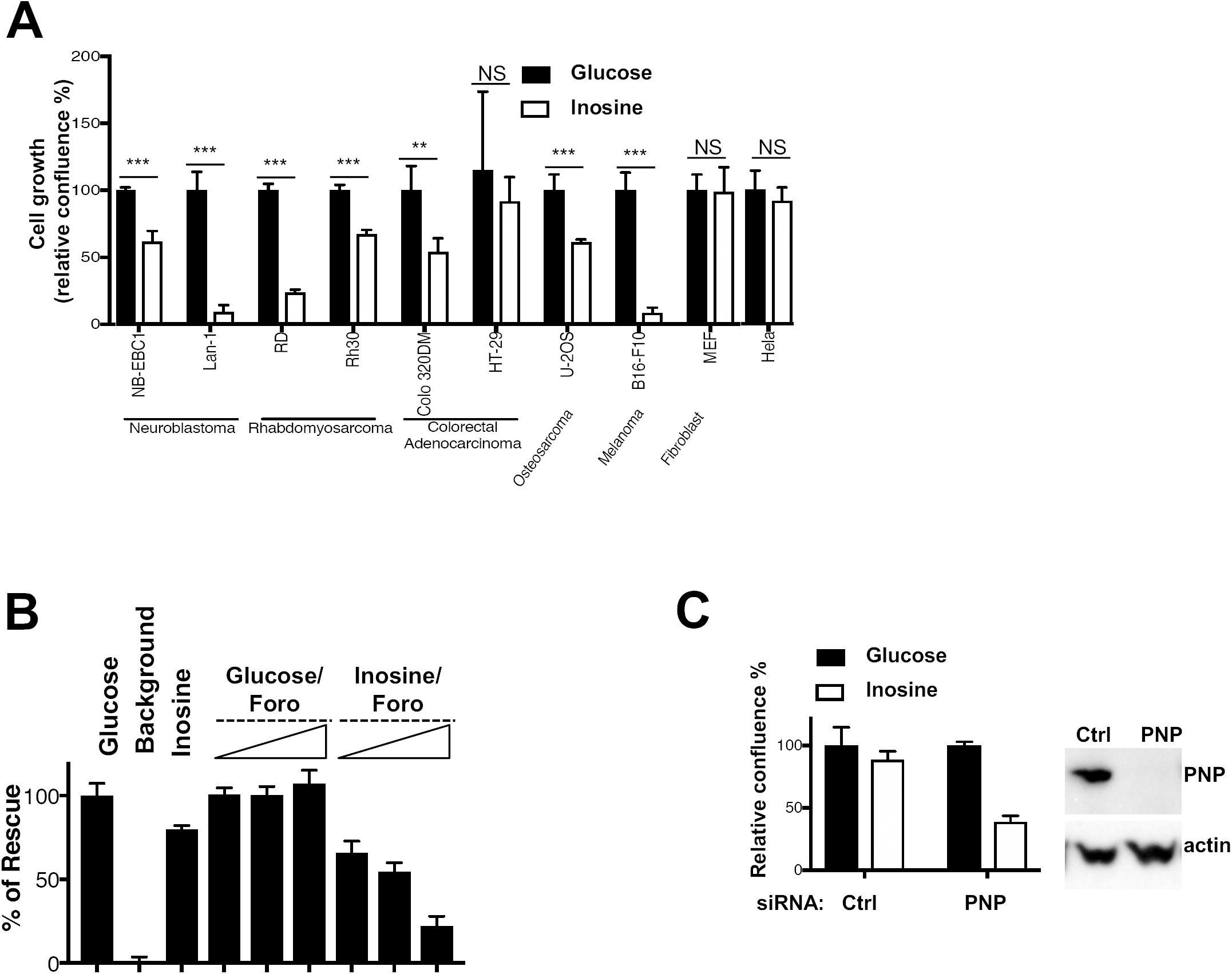
Transformed cells display diverse capacity to utilize inosine as a carbon source. (A) The indicated cell lines were cultured without glucose (background), with glucose or with inosine, and cell growth curves were monitored by live cell imaging analysis (IncuCyte ZOOM™). Cell confluences of the indicated cell lines cultured with glucose at 72-96 h were set to 100%. Error bars represent mean ± SD (n=4). **, *P*<0.01 and ***, *P*<0.001 for Glucose versus Inosine, respectively. (B) HeLa cells were incubated without glucose (background), with glucose or with inosine, as well as in combination with increased concentrations of Foro (0.1 μM, 0.5 μM, and 2 μM), and cell growth was monitored by live cell imaging analysis (IncuCyte ZOOM™). The confluences of HeLa cells cultured with glucose at 60 h were set to 100%. Error bars represent mean ± SD (n=4). Data are representative of two independent experiments. (C) HeLa cells were transfected with scrambled siRNA or PNP siRNA for 3 days. Cells were then switched to media without glucose (background), with either glucose or inosine, and cell confluence was determined by live cell imaging analysis (IncuCyte ZOOM™). The confluences of HeLa cells in the presence of glucose at 96 h were set to 100%. Error bars represent standard deviation from mean of quadruplicate samples (left). PNP protein expression levels were determined by immunoblot (right). Data are representative of two independent experiments.

**Figure S7.**
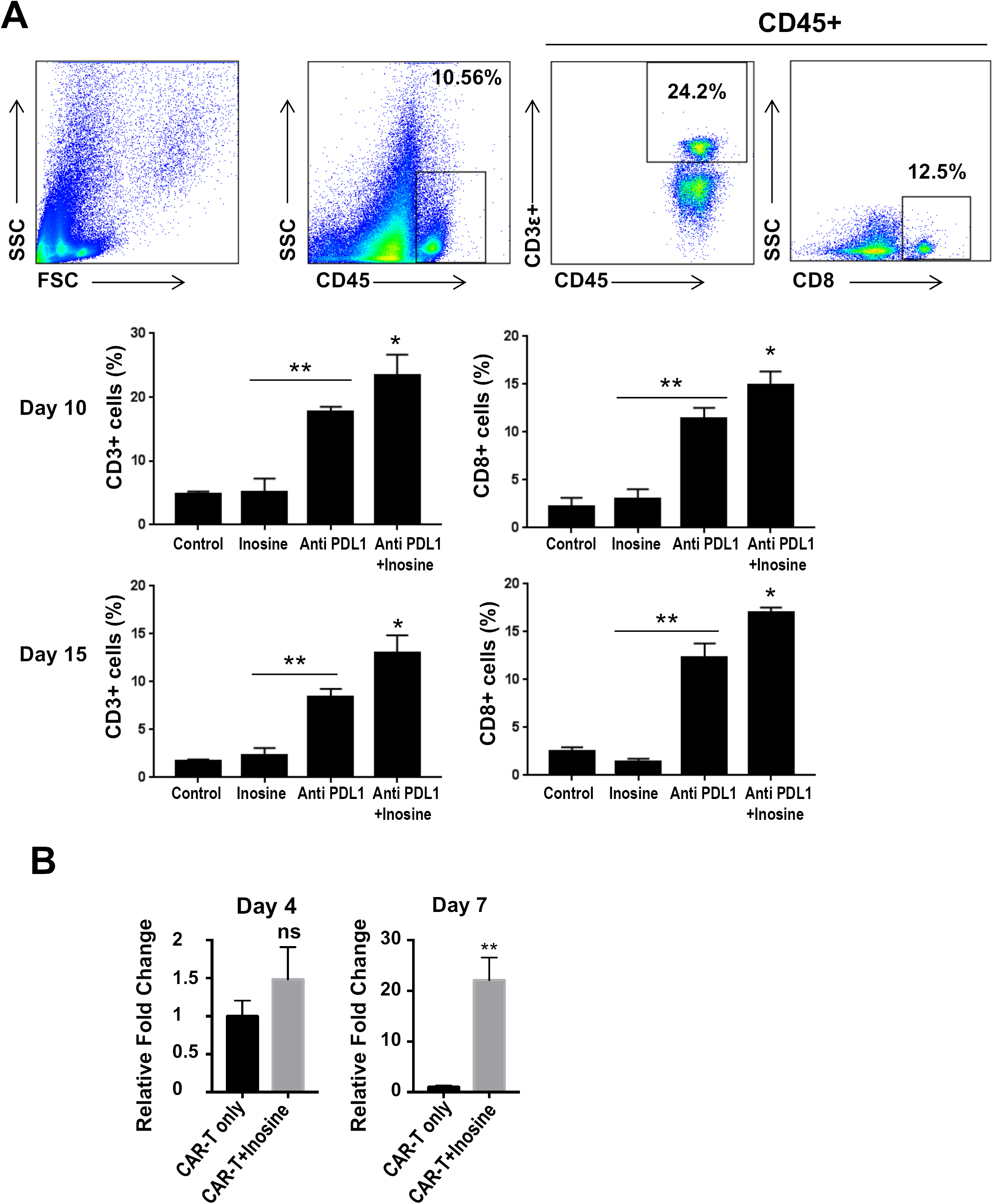
Inosine supplement prolongs CAR-T cell proliferation and survival in tumor environment. (A) Murine melanoma xenograft model was established in C57BL/6 mice by subcutaneously inoculation of B16F10 tumor cells. T cell infiltration in tumor was monitored at indicated time by flow cytometry after surface staining of anti-CD45, anti-CD3 and anti-CD8 antibodies. Gating strategy for flow cytometry analysis of leucocytes (CD45+), T cells (CD3+), and effector T cells (CD3+CD8+) (top panel); Values represent mean ± SEM (n=3). *, *P*< 0.05 for anti-PDL1 versus anti-PDL1+ inosine and **, *P*< 0.001 for Control versus anti-PDL1. (B) Human neuroblastoma xenograft model was established in NSG mice by subcutaneously inoculation of LAN-1 neuroblastoma cells, followed by i.v. administration of GD2-CAR-T-Luciferase cells only or in combination with oral gavage of inosine (300 mg/kg/day) when tumor grow to 4-6 mm in diameter. T cell infiltration in tumor was monitored at indicated time by IVIS imaging. Error bars represent mean ± SEM (n=5). **, *P*< 0.01 for CAR-T only versus CAR-T + inosine at Day 7. Data are representative of two independent experiments.

## References

1. D. Finlay, and D.A. Cantrell, Metabolism, migration and memory in cytotoxic T cells. Nature reviews. Immunology 11 (2011) 109–17.

2. R. Wang, and D.R. Green, Metabolic checkpoints in activated T cells. Nature immunology 13 (2012) 907–15.

3. E.L. Pearce, and E.J. Pearce, Metabolic pathways in immune cell activation and quiescence. Immunity 38 (2013) 633–43.

4. S.E. Weinberg, L.A. Sena, and N.S. Chandel, Mitochondria in the regulation of innate and adaptive immunity. Immunity 42 (2015) 406–17.

5. L.A. O’Neill, R.J. Kishton, and J. Rathmell, A guide to immunometabolism for immunologists. Nature reviews. Immunology 16 (2016) 553–65.

6. M.D. Buck, R.T. Sowell, S.M. Kaech, and E.L. Pearce, Metabolic Instruction of Immunity. Cell 169 (2017) 570–586.

7. E.H. Ma, M.C. Poffenberger, A.H. Wong, and R.G. Jones, The role of AMPK in T cell metabolism and function. Current opinion in immunology 46 (2017) 45–52.

8. C.H. Patel, and J.D. Powell, Targeting T cell metabolism to regulate T cell activation, differentiation and function in disease. Current opinion in immunology 46 (2017) 82–88.

9. H. Zeng, and H. Chi, mTOR signaling in the differentiation and function of regulatory and effector T cells. Current opinion in immunology 46 (2017) 103–111.

10. G. Lian, J.R. Gnanaprakasam, T. Wang, R. Wu, X. Chen, L. Liu, Y. Shen, M. Yang, J. Yang, Y. Chen, V. Vasiliou, T.A. Cassel, D.R. Green, Y. Liu, T.W. Fan, and R. Wang, Glutathione de novo synthesis but not recycling process coordinates with glutamine catabolism to control redox homeostasis and directs murine T cell differentiation. eLife 7 (2018).

11. M. Slack, T. Wang, and R. Wang, T cell metabolic reprogramming and plasticity. Molecular immunology 68 (2015) 507–12.

12. J.C. Rathmell, E.A. Farkash, W. Gao, and C.B. Thompson, IL-7 enhances the survival and maintains the size of naive T cells. J Immunol 167 (2001) 6869–76.

13. K.A. Frauwirth, J.L. Riley, M.H. Harris, R.V. Parry, J.C. Rathmell, D.R. Plas, R.L. Elstrom, C.H. June, and C.B. Thompson, The CD28 signaling pathway regulates glucose metabolism. Immunity 16 (2002) 769–77.

14. S.R. Jacobs, C.E. Herman, N.J. Maciver, J.A. Wofford, H.L. Wieman, J.J. Hammen, and J.C. Rathmell, Glucose uptake is limiting in T cell activation and requires CD28-mediated Akt-dependent and independent pathways. J Immunol 180 (2008) 4476–86.

15. E.L. Pearce, M.C. Walsh, P.J. Cejas, G.M. Harms, H. Shen, L.S. Wang, R.G. Jones, and Y. Choi, Enhancing CD8 T-cell memory by modulating fatty acid metabolism. Nature 460 (2009) 103–7.

16. R. Wang, C.P. Dillon, L.Z. Shi, S. Milasta, R. Carter, D. Finkelstein, L.L. McCormick, P. Fitzgerald, H. Chi, J. Munger, and D.R. Green, The Transcription Factor Myc Controls Metabolic Reprogramming upon T Lymphocyte Activation. Immunity 35 (2011) 871–82.

17. V.A. Gerriets, and J.C. Rathmell, Metabolic pathways in T cell fate and function. Trends Immunol 33 (2012) 168–73.

18. G.J. van der Windt, B. Everts, C.H. Chang, J.D. Curtis, T.C. Freitas, E. Amiel, E.J. Pearce, and E.L. Pearce, Mitochondrial Respiratory Capacity Is a Critical Regulator of CD8(+) T Cell Memory Development. Immunity (2012).

19. M. Angela, Y. Endo, H.K. Asou, T. Yamamoto, D.J. Tumes, H. Tokuyama, K. Yokote, and T. Nakayama, Fatty acid metabolic reprogramming via mTOR-mediated inductions of PPARgamma directs early activation of T cells. Nature communications 7 (2016) 13683.

20. M.D. Buck, D. O’Sullivan, R.I. Klein Geltink, J.D. Curtis, C.H. Chang, D.E. Sanin, J. Qiu, O. Kretz, D. Braas, G.J. van der Windt, Q. Chen, S.C. Huang, C.M. O’Neill, B.T. Edelson, E.J. Pearce, H. Sesaki, T.B. Huber, A.S. Rambold, and E.L. Pearce, Mitochondrial Dynamics Controls T Cell Fate through Metabolic Programming. Cell 166 (2016) 63–76.

21. S.J. Bensinger, M.N. Bradley, S.B. Joseph, N. Zelcer, E.M. Janssen, M.A. Hausner, R. Shih, J.S. Parks, P.A. Edwards, B.D. Jamieson, and P. Tontonoz, LXR signaling couples sterol metabolism to proliferation in the acquired immune response. Cell 134 (2008) 97–111.

22. J.A. Wofford, H.L. Wieman, S.R. Jacobs, Y. Zhao, and J.C. Rathmell, IL-7 promotes Glut1 trafficking and glucose uptake via STAT5-mediated activation of Akt to support T-cell survival. Blood 111 (2008) 2101–11.

23. E.L. Carr, A. Kelman, G.S. Wu, R. Gopaul, E. Senkevitch, A. Aghvanyan, A.M. Turay, and K.A. Frauwirth, Glutamine uptake and metabolism are coordinately regulated by ERK/MAPK during T lymphocyte activation. J Immunol 185 (2011) 1037–44.

24. R.D. Michalek, V.A. Gerriets, S.R. Jacobs, A.N. Macintyre, N.J. MacIver, E.F. Mason, S.A. Sullivan, A.G. Nichols, and J.C. Rathmell, Cutting edge: Distinct glycolytic and lipid oxidative metabolic programs are essential for effector and regulatory CD4+ T cell subsets. J Immunol 186 (2011) 3299–303.

25. D.K. Finlay, E. Rosenzweig, L.V. Sinclair, C. Feijoo-Carnero, J.L. Hukelmann, J. Rolf, A.A. Panteleyev, K. Okkenhaug, and D.A. Cantrell, PDK1 regulation of mTOR and hypoxia-inducible factor 1 integrate metabolism and migration of CD8+ T cells. The Journal of experimental medicine 209 (2012) 2441–53.

26. Y. Kidani, H. Elsaesser, M.B. Hock, L. Vergnes, K.J. Williams, J.P. Argus, B.N. Marbois, E. Komisopoulou, E.B. Wilson, T.F. Osborne, T.G. Graeber, K. Reue, D.G. Brooks, and S.J. Bensinger, Sterol regulatory element-binding proteins are essential for the metabolic programming of effector T cells and adaptive immunity. Nature immunology 14 (2013) 489–99.

27. S.I. Grivennikov, F.R. Greten, and M. Karin, Immunity, inflammation, and cancer. Cell 140 (2010) 883–99.

28. R.D. Schreiber, L.J. Old, and M.J. Smyth, Cancer immunoediting: integrating immunity’s roles in cancer suppression and promotion. Science 331 (2011) 1565–70.

29. W.H. Fridman, F. Pages, C. Sautes-Fridman, and J. Galon, The immune contexture in human tumours: impact on clinical outcome. Nature reviews. Cancer 12 (2012) 298–306.

30. N.P. Restifo, M.E. Dudley, and S.A. Rosenberg, Adoptive immunotherapy for cancer: harnessing the T cell response. Nature reviews. Immunology 12 (2012) 269–81.

31. T.F. Gajewski, H. Schreiber, and Y.X. Fu, Innate and adaptive immune cells in the tumor microenvironment. Nature immunology 14 (2013) 1014–22.

32. A.K. Palucka, and L.M. Coussens, The Basis of Oncoimmunology. Cell 164 (2016) 1233–47.

33. D. Mittal, M.M. Gubin, R.D. Schreiber, and M.J. Smyth, New insights into cancer immunoediting and its three component phases--elimination, equilibrium and escape. Current opinion in immunology 27 (2014) 16–25.

34. R. Roychoudhuri, R.L. Eil, and N.P. Restifo, The interplay of effector and regulatory T cells in cancer. Current opinion in immunology 33 (2015) 101–11.

35. D.E. Speiser, P.C. Ho, and G. Verdeil, Regulatory circuits of T cell function in cancer. Nature reviews. Immunology 16 (2016) 599–611.

36. H. Nishikawa, and S. Sakaguchi, Regulatory T cells in cancer immunotherapy. Current opinion in immunology 27 (2014) 1–7.

37. P. Sharma, and J.P. Allison, Immune checkpoint targeting in cancer therapy: toward combination strategies with curative potential. Cell 161 (2015) 205–14.

38. C. Engblom, C. Pfirschke, and M.J. Pittet, The role of myeloid cells in cancer therapies. Nature reviews. Cancer 16 (2016) 447–62.

39. V. Kumar, S. Patel, E. Tcyganov, and D.I. Gabrilovich, The Nature of Myeloid-Derived Suppressor Cells in the Tumor Microenvironment. Trends Immunol 37 (2016) 208–220.

40. W. Zou, J.D. Wolchok, and L. Chen, PD-L1 (B7-H1) and PD-1 pathway blockade for cancer therapy: Mechanisms, response biomarkers, and combinations. Science translational medicine 8 (2016) 328rv4.

41. L.P. Andrews, A.E. Marciscano, C.G. Drake, and D.A. Vignali, LAG3 (CD223) as a cancer immunotherapy target. Immunological reviews 276 (2017) 80–96.

42. D.M. Barrett, N. Singh, D.L. Porter, S.A. Grupp, and C.H. June, Chimeric antigen receptor therapy for cancer. Annual review of medicine 65 (2014) 333–47.

43. L. Chen, and X. Han, Anti-PD-1/PD-L1 therapy of human cancer: past, present, and future. The Journal of clinical investigation 125 (2015) 3384–91.

44. S.A. Rosenberg, and N.P. Restifo, Adoptive cell transfer as personalized immunotherapy for human cancer. Science 348 (2015) 62–8.

45. P. Sharma, and J.P. Allison, The future of immune checkpoint therapy. Science 348 (2015) 56–61.

46. A.D. Fesnak, C.H. June, and B.L. Levine, Engineered T cells: the promise and challenges of cancer immunotherapy. Nature reviews. Cancer 16 (2016) 566–81.

47. H.J. Jackson, S. Rafiq, and R.J. Brentjens, Driving CAR T-cells forward. Nature reviews. Clinical oncology 13 (2016) 370–83.

48. A.J. Minn, and E.J. Wherry, Combination Cancer Therapies with Immune Checkpoint Blockade: Convergence on Interferon Signaling. Cell 165 (2016) 272–5.

49. W.A. Lim, and C.H. June, The Principles of Engineering Immune Cells to Treat Cancer. Cell 168 (2017) 724–740.

50. D.E. Gilham, R. Debets, M. Pule, R.E. Hawkins, and H. Abken, CAR-T cells and solid tumors: tuning T cells to challenge an inveterate foe. Trends in molecular medicine 18 (2012) 377–84.

51. M. Sadelain, I. Riviere, and S. Riddell, Therapeutic T cell engineering. Nature 545 (2017) 423–431.

52. P. Sharma, S. Hu-Lieskovan, J.A. Wargo, and A. Ribas, Primary, Adaptive, and Acquired Resistance to Cancer Immunotherapy. Cell 168 (2017) 707–723.

53. J.J. Havel, D. Chowell, and T.A. Chan, The evolving landscape of biomarkers for checkpoint inhibitor immunotherapy. Nature reviews. Cancer 19 (2019) 133–150.

54. V.M. Konala, S. Adapa, and W.S. Aronow, Immunotherapy in Bladder Cancer. American journal of therapeutics (2019).

55. P.P. Hsu, and D.M. Sabatini, Cancer cell metabolism: Warburg and beyond. Cell 134 (2008) 703–7.

56. M.G. Vander Heiden, L.C. Cantley, and C.B. Thompson, Understanding the Warburg effect: the metabolic requirements of cell proliferation. Science 324 (2009) 1029–33.

57. R.A. Cairns, I.S. Harris, and T.W. Mak, Regulation of cancer cell metabolism. Nature reviews. Cancer 11 (2011) 85–95.

58. D. Hanahan, and R.A. Weinberg, Hallmarks of cancer: the next generation. Cell 144 (2011) 646–74.

59. P.S. Ward, and C.B. Thompson, Metabolic reprogramming: a cancer hallmark even warburg did not anticipate. Cancer cell 21 (2012) 297–308.

60. C.V. Dang, MYC, metabolism, cell growth, and tumorigenesis. Cold Spring Harbor perspectives in medicine 3 (2013).

61. G.M. DeNicola, and L.C. Cantley, Cancer’s Fuel Choice: New Flavors for a Picky Eater. Molecular cell 60 (2015) 514–23.

62. D. O’Sullivan, and E.L. Pearce, Targeting T cell metabolism for therapy. Trends Immunol 36 (2015) 71–80.

63. N.N. Pavlova, and C.B. Thompson, The Emerging Hallmarks of Cancer Metabolism. Cell metabolism 23 (2016) 27–47.

64. C.M. Cham, and T.F. Gajewski, Glucose availability regulates IFN-gamma production and p70S6 kinase activation in CD8+ effector T cells. Journal of immunology 174 (2005) 4670–7.

65. L. Antonioli, C. Blandizzi, P. Pacher, and G. Hasko, Immunity, inflammation and cancer: a leading role for adenosine. Nature reviews. Cancer 13 (2013) 842–57.

66. T. Wang, G. Liu, and R. Wang, The Intercellular Metabolic Interplay between Tumor and Immune Cells. Frontiers in immunology 5 (2014) 358.

67. C.H. Chang, J. Qiu, D. O’Sullivan, M.D. Buck, T. Noguchi, J.D. Curtis, Q. Chen, M. Gindin, M.M. Gubin, G.J. van der Windt, E. Tonc, R.D. Schreiber, E.J. Pearce, and E.L. Pearce, Metabolic Competition in the Tumor Microenvironment Is a Driver of Cancer Progression. Cell 162 (2015) 1229–41.

68. P.C. Ho, J.D. Bihuniak, A.N. Macintyre, M. Staron, X. Liu, R. Amezquita, Y.C. Tsui, G. Cui, G. Micevic, J.C. Perales, S.H. Kleinstein, E.D. Abel, K.L. Insogna, S. Feske, J.W. Locasale, M.W. Bosenberg, J.C. Rathmell, and S.M. Kaech, Phosphoenolpyruvate Is a Metabolic Checkpoint of Anti-tumor T Cell Responses. Cell 162 (2015) 1217–28.

69. S. Kleffel, C. Posch, S.R. Barthel, H. Mueller, C. Schlapbach, E. Guenova, C.P. Elco, N. Lee, V.R. Juneja, Q. Zhan, C.G. Lian, R. Thomi, W. Hoetzenecker, A. Cozzio, R. Dummer, M.C. Mihm, Jr., K.T. Flaherty, M.H. Frank, G.F. Murphy, A.H. Sharpe, T.S. Kupper, and T. Schatton, Melanoma Cell-Intrinsic PD-1 Receptor Functions Promote Tumor Growth. Cell 162 (2015) 1242–56.

70. C. Herbel, N. Patsoukis, K. Bardhan, P. Seth, J.D. Weaver, and V.A. Boussiotis, Clinical significance of T cell metabolic reprogramming in cancer. Clinical and translational medicine 5 (2016) 29.

71. D.H. Munn, and A.L. Mellor, IDO in the Tumor Microenvironment: Inflammation, Counter-Regulation, and Tolerance. Trends Immunol 37 (2016) 193–207.

72. G. Andrejeva, and J.C. Rathmell, Similarities and Distinctions of Cancer and Immune Metabolism in Inflammation and Tumors. Cell metabolism 26 (2017) 49–70.

73. R.J. Kishton, M. Sukumar, and N.P. Restifo, Metabolic Regulation of T Cell Longevity and Function in Tumor Immunotherapy. Cell metabolism 26 (2017) 94–109.

74. X. Xu, J.N.R. Gnanaprakasam, J. Sherman, and R. Wang, A Metabolism Toolbox for CAR T Therapy. Frontiers in oncology 9 (2019) 322.

75. L.Z. Shi, R. Wang, G. Huang, P. Vogel, G. Neale, D.R. Green, and H. Chi, HIF1alpha-dependent glycolytic pathway orchestrates a metabolic checkpoint for the differentiation of TH17 and Treg cells. The Journal of experimental medicine 208 (2011) 1367–76.

76. B.R. Bochner, M. Siri, R.H. Huang, S. Noble, X.H. Lei, P.A. Clemons, and B.K. Wagner, Assay of the multiple energy-producing pathways of mammalian cells. PLoS One 6 (2011) e18147.

77. W.W. Overwijk, A. Tsung, K.R. Irvine, M.R. Parkhurst, T.J. Goletz, K. Tsung, M.W. Carroll, C. Liu, B. Moss, S.A. Rosenberg, and N.P. Restifo, gp100/pmel 17 is a murine tumor rejection antigen: induction of “self”-reactive, tumoricidal T cells using high-affinity, altered peptide ligand. The Journal of experimental medicine 188 (1998) 277–86.

78. W.W. Overwijk, D.S. Lee, D.R. Surman, K.R. Irvine, C.E. Touloukian, C.C. Chan, M.W. Carroll, B. Moss, S.A. Rosenberg, and N.P. Restifo, Vaccination with a recombinant vaccinia virus encoding a “self” antigen induces autoimmune vitiligo and tumor cell destruction in mice: requirement for CD4(+) T lymphocytes. Proceedings of the National Academy of Sciences of the United States of America 96 (1999) 2982–7.

79. W.W. Overwijk, M.R. Theoret, S.E. Finkelstein, D.R. Surman, L.A. de Jong, F.A. Vyth- Dreese, T.A. Dellemijn, P.A. Antony, P.J. Spiess, D.C. Palmer, D.M. Heimann, C.A. Klebanoff, Z. Yu, L.N. Hwang, L. Feigenbaum, A.M. Kruisbeek, S.A. Rosenberg, and N.P. Restifo, Tumor regression and autoimmunity after reversal of a functionally tolerant state of self-reactive CD8+ T cells. The Journal of experimental medicine 198 (2003) 569–80.

80. M.A. Pule, B. Savoldo, G.D. Myers, C. Rossig, H.V. Russell, G. Dotti, M.H. Huls, E. Liu, A.P. Gee, Z. Mei, E. Yvon, H.L. Weiss, H. Liu, C.M. Rooney, H.E. Heslop, and M.K. Brenner, Virus-specific T cells engineered to coexpress tumor-specific receptors: persistence and antitumor activity in individuals with neuroblastoma. Nature medicine 14 (2008) 1264–70.

81. J.A. Craddock, A. Lu, A. Bear, M. Pule, M.K. Brenner, C.M. Rooney, and A.E. Foster, Enhanced tumor trafficking of GD2 chimeric antigen receptor T cells by expression of the chemokine receptor CCR2b. Journal of immunotherapy 33 (2010) 780–8.

82. G. Burnstock, Purinergic signalling--an overview. Novartis Foundation symposium 276 (2006) 26-48; discussion 48-57, 275-81.

83. G. Burnstock, Purinergic signalling and disorders of the central nervous system. Nature reviews. Drug discovery 7 (2008) 575–90.

84. H.K. Eltzschig, M.V. Sitkovsky, and S.C. Robson, Purinergic signaling during inflammation. The New England journal of medicine 367 (2012) 2322–33.

85. C. Cekic, and J. Linden, Purinergic regulation of the immune system. Nature reviews. Immunology 16 (2016) 177–92.

86. J.N. Rashida Gnanaprakasam, R. Wu, and R. Wang, Metabolic Reprogramming in Modulating T Cell Reactive Oxygen Species Generation and Antioxidant Capacity. Frontiers in immunology 9 (2018) 1075.

87. A. Bzowska, E. Kulikowska, and D. Shugar, Purine nucleoside phosphorylases: properties, functions, and clinical aspects. Pharmacology & therapeutics 88 (2000) 349–425.

88. R.G. Silva, J.E. Nunes, F. Canduri, J.C. Borges, L.M. Gava, F.B. Moreno, L.A. Basso, and D.S. Santos, Purine nucleoside phosphorylase: a potential target for the development of drugs to treat T-cell- and apicomplexan parasite-mediated diseases. Current drug targets 8 (2007) 413–22.

89. A. Korycka, J.Z. Blonski, and T. Robak, Forodesine (BCX-1777, Immucillin H)--a new purine nucleoside analogue: mechanism of action and potential clinical application. Mini reviews in medicinal chemistry 7 (2007) 976–83.

90. A. Al-Kali, V. Gandhi, M. Ayoubi, M. Keating, and F. Ravandi, Forodesine: review of preclinical and clinical data. Future oncology 6 (2010) 1211–7.

91. A. Kratz, M. Ferraro, P.M. Sluss, and K.B. Lewandrowski, Case records of the Massachusetts General Hospital. Weekly clinicopathological exercises. Laboratory reference values. The New England journal of medicine 351 (2004) 1548–63.

92. N. Psychogios, D.D. Hau, J. Peng, A.C. Guo, R. Mandal, S. Bouatra, I. Sinelnikov, R. Krishnamurthy, R. Eisner, B. Gautam, N. Young, J. Xia, C. Knox, E. Dong, P. Huang, Z. Hollander, T.L. Pedersen, S.R. Smith, F. Bamforth, R. Greiner, B. McManus, J.W. Newman, T. Goodfriend, and D.S. Wishart, The human serum metabolome. PLoS One 6 (2011) e16957.

93. J.R. Mayers, and M.G. Vander Heiden, Famine versus feast: understanding the metabolism of tumors in vivo. Trends in biochemical sciences 40 (2015) 130–40.

94. C. Commisso, S.M. Davidson, R.G. Soydaner-Azeloglu, S.J. Parker, J.J. Kamphorst, S. Hackett, E. Grabocka, M. Nofal, J.A. Drebin, C.B. Thompson, J.D. Rabinowitz, C.M. Metallo, M.G. Vander Heiden, and D. Bar-Sagi, Macropinocytosis of protein is an amino acid supply route in Ras-transformed cells. Nature 497 (2013) 633–7.

95. T. Mashimo, K. Pichumani, V. Vemireddy, K.J. Hatanpaa, D.K. Singh, S. Sirasanagandla, S. Nannepaga, S.G. Piccirillo, Z. Kovacs, C. Foong, Z. Huang, S. Barnett, B.E. Mickey, R.J. DeBerardinis, B.P. Tu, E.A. Maher, and R.M. Bachoo, Acetate is a bioenergetic substrate for human glioblastoma and brain metastases. Cell 159 (2014) 1603–14.

96. S.K. Shukla, T. Gebregiworgis, V. Purohit, N.V. Chaika, V. Gunda, P. Radhakrishnan, K. Mehla, Pipinos, II, R. Powers, F. Yu, and P.K. Singh, Metabolic reprogramming induced by ketone bodies diminishes pancreatic cancer cachexia. Cancer & metabolism 2 (2014) 18.

97. C.H. Chang, J.D. Curtis, L.B. Maggi, Jr., B. Faubert, A.V. Villarino, D. O’Sullivan, S.C. Huang, G.J. van der Windt, J. Blagih, J. Qiu, J.D. Weber, E.J. Pearce, R.G. Jones, and E.L. Pearce, Posttranscriptional control of T cell effector function by aerobic glycolysis. Cell 153 (2013) 1239–51.

98. E. Kun, P. Talalay, and H.G. Williams-Ashman, Studies on the Ehrlich ascites tumor. I. The enzymic and metabolic activities of the ascitic cells and the ascitic plasma. Cancer research 11 (1951) 855–63.

99. H. Harrington, Effect of glucose and various nucleosides on purine synthesis by Ehrlich ascites tumor cells in vitro. The Journal of biological chemistry 233 (1958) 1190–3.

100. M. Cappiello, D. Barsacchi, A. Del Corso, M.G. Tozzi, M. Camici, U. Mura, and P.L. Ipata, Purine salvage as a metabolite and energy saving mechanism in the ocular lens. Current eye research 11 (1992) 435–44.

101. Y.F. Xu, F. Letisse, F. Absalan, W. Lu, E. Kuznetsova, G. Brown, A.A. Caudy, A.F. Yakunin, J.R. Broach, and J.D. Rabinowitz, Nucleotide degradation and ribose salvage in yeast. Molecular systems biology 9 (2013) 665.

102. S. Tabata, M. Yamamoto, H. Goto, A. Hirayama, M. Ohishi, T. Kuramoto, A. Mitsuhashi, R. Ikeda, M. Haraguchi, K. Kawahara, Y. Shinsato, K. Minami, A. Saijo, M. Hanibuchi, Y. Nishioka, S. Sone, H. Esumi, M. Tomita, T. Soga, T. Furukawa, and S.I. Akiyama, Thymidine Catabolism as a Metabolic Strategy for Cancer Survival. Cell reports 19 (2017) 1313–1321.

103. A.L. Koch, Some enzymes of nucleoside metabolism of Escherichia coli. The Journal of biological chemistry 223 (1956) 535–49.

104. K. Hammer-Jespersen, Nucleoside catabolism. In Metabolism of Nucleotides. in: A. Munch-Petersen, (Ed.), Nucleosides and Nucleobases in Microorganisms, Academic Press, New York, 1983, pp. 203–258.

105. R. Schuch, A. Garibian, H.H. Saxild, P.J. Piggot, and P. Nygaard, Nucleosides as a carbon source in Bacillus subtilis: characterization of the drm-pupG operon. Microbiology 145 (Pt 10) (1999) 2957–66.

106. M.G. Tozzi, M. Camici, L. Mascia, F. Sgarrella, and P.L. Ipata, Pentose phosphates in nucleoside interconversion and catabolism. The FEBS journal 273 (2006) 1089–101.

107. T. Rimaux, G. Vrancken, B. Vuylsteke, L. De Vuyst, and F. Leroy, The pentose moiety of adenosine and inosine is an important energy source for the fermented-meat starter culture Lactobacillus sakei CTC 494. Applied and environmental microbiology 77 (2011) 6539–50.

108. R.D. Lange, W.H. Crosby, D.M. Donohue, C.A. Finch, J.G. Gibson, 2nd, M.T. Mc, and M.M. Strumia, Effect of inosine on red cell preservation. The Journal of clinical investigation 37 (1958) 1485–93.

109. A.W. Konings, Comparison of inosine and glucose as a substrate for energy metabolism in isolated rat-thymus nuclei. Biochimica et biophysica acta 189 (1969) 125–8.

110. M.M. Strumia, and P.V. Strumia, The preservation of blood for transfusion. IX. The effect of increased pH and addition of inosine only or adenine and inosine on the red cell function. The Journal of laboratory and clinical medicine 79 (1972) 863–72.

111. A.R. Fernando, D.M. Armstrong, J.R. Griffiths, W.F. Hendry, E.P. O’Donoghue, D. Perrett, J.P. Ward, and J.E. Wickham, Enhanced preservation of the ischaemic kidney with inosine. Lancet 1 (1976) 555–7.

112. S.M. Jarvis, J.D. Young, M. Ansay, A.L. Archibald, R.A. Harkness, and R.J. Simmonds, Is inosine the physiological energy source of pig erythrocytes? Biochimica et biophysica acta 597 (1980) 183–8.

113. R.B. Jennings, and C. Steenbergen, Jr., Nucleotide metabolism and cellular damage in myocardial ischemia. Annual review of physiology 47 (1985) 727–49.

114. M.D. Devous, Sr., and E.D. Lewandowski, Inosine preserves ATP during ischemia and enhances recovery during reperfusion. The American journal of physiology 253 (1987) H1224–33.

115. A. Mathew, M. Grdisa, and R.M. Johnstone, Nucleosides and glutamine are primary energy substrates for embryonic and adult chicken red cells. Biochemistry and cell biology = Biochimie et biologie cellulaire 71 (1993) 288–95.

116. M.S. Jurkowitz, M.L. Litsky, M.J. Browning, and C.M. Hohl, Adenosine, inosine, and guanosine protect glial cells during glucose deprivation and mitochondrial inhibition: correlation between protection and ATP preservation. Journal of neurochemistry 71 (1998) 535–48.

117. S.E. Haun, J.E. Segeleon, V.L. Trapp, M.A. Clotz, and L.A. Horrocks, Inosine mediates the protective effect of adenosine in rat astrocyte cultures subjected to combined glucose-oxygen deprivation. Journal of neurochemistry 67 (1996) 2051–9.

118. T.W. Traut, Physiological concentrations of purines and pyrimidines. Molecular and cellular biochemistry 140 (1994) 1–22.

119. I. Kaufmann, A. Hoelzl, F. Schliephake, T. Hummel, A. Chouker, L. Lysenko, K. Peter, and M. Thiel, Effects of adenosine on functions of polymorphonuclear leukocytes from patients with septic shock. Shock (Augusta, Ga.) 27 (2007) 25–31.

120. P. Pellegatti, L. Raffaghello, G. Bianchi, F. Piccardi, V. Pistoia, and F. Di Virgilio, Increased level of extracellular ATP at tumor sites: in vivo imaging with plasma membrane luciferase. PLoS One 3 (2008) e2599.

121. J. Stagg, and M.J. Smyth, Extracellular adenosine triphosphate and adenosine in cancer. Oncogene 29 (2010) 5346–58.

122. P.L. Ipata, M. Camici, V. Micheli, and M.G. Tozz, Metabolic network of nucleosides in the brain. Current topics in medicinal chemistry 11 (2011) 909–22.

123. V. Kumar, Adenosine as an endogenous immunoregulator in cancer pathogenesis: where to go? Purinergic signalling 9 (2013) 145–65.

124. M.P. Abbracchio, G. Burnstock, A. Verkhratsky, and H. Zimmermann, Purinergic signalling in the nervous system: an overview. Trends in neurosciences 32 (2009) 19–29.

125. W.G. Junger, Immune cell regulation by autocrine purinergic signalling. Nature reviews. Immunology 11 (2011) 201–12.

126. X. Jin, R.K. Shepherd, B.R. Duling, and J. Linden, Inosine binds to A3 adenosine receptors and stimulates mast cell degranulation. The Journal of clinical investigation 100 (1997) 2849–57.

127. E.R. Giblett, J.E. Anderson, F. Cohen, B. Pollara, and H.J. Meuwissen, Adenosine-deaminase deficiency in two patients with severely impaired cellular immunity. Lancet 2 (1972) 1067–9.

128. R. Parkman, E.W. Gelfand, F.S. Rosen, A. Sanderson, and R. Hirschhorn, Severe combined immunodeficiency and adenosine deaminase deficiency. The New England journal of medicine 292 (1975) 714–9.

129. M.S. Hershfield, R.H. Buckley, M.L. Greenberg, A.L. Melton, R. Schiff, C. Hatem, J. Kurtzberg, M.L. Markert, R.H. Kobayashi, A.L. Kobayashi, and et al., Treatment of adenosine deaminase deficiency with polyethylene glycol-modified adenosine deaminase. The New England journal of medicine 316 (1987) 589–96.

130. A. Aiuti, F. Cattaneo, S. Galimberti, U. Benninghoff, B. Cassani, L. Callegaro, S. Scaramuzza, G. Andolfi, M. Mirolo, I. Brigida, A. Tabucchi, F. Carlucci, M. Eibl, M. Aker, S. Slavin, H. Al-Mousa, A. Al Ghonaium, A. Ferster, A. Duppenthaler, L. Notarangelo, U. Wintergerst, R.H. Buckley, M. Bregni, S. Marktel, M.G. Valsecchi, P. Rossi, F. Ciceri, R. Miniero, C. Bordignon, and M.G. Roncarolo, Gene therapy for immunodeficiency due to adenosine deaminase deficiency. The New England journal of medicine 360 (2009) 447–58.

131. M. Sukumar, J. Liu, Y. Ji, M. Subramanian, J.G. Crompton, Z. Yu, R. Roychoudhuri, D.C. Palmer, P. Muranski, E.D. Karoly, R.P. Mohney, C.A. Klebanoff, A. Lal, T. Finkel, N.P. Restifo, and L. Gattinoni, Inhibiting glycolytic metabolism enhances CD8+ T cell memory and antitumor function. The Journal of clinical investigation 123 (2013) 4479–88.

132. A. Caro-Maldonado, R. Wang, A.G. Nichols, M. Kuraoka, S. Milasta, L.D. Sun, A.L. Gavin, E.D. Abel, G. Kelsoe, D.R. Green, and J.C. Rathmell, Metabolic reprogramming is required for antibody production that is suppressed in anergic but exaggerated in chronically BAFF-exposed B cells. J Immunol 192 (2014) 3626–36.

133. V.A. Gerriets, R.J. Kishton, A.G. Nichols, A.N. Macintyre, M. Inoue, O. Ilkayeva, P.S. Winter, X. Liu, B. Priyadharshini, M.E. Slawinska, L. Haeberli, C. Huck, L.A. Turka, K.C. Wood, L.P. Hale, P.A. Smith, M.A. Schneider, N.J. MacIver, J.W. Locasale, C.B. Newgard, M.L. Shinohara, and J.C. Rathmell, Metabolic programming and PDHK1 control CD4+ T cell subsets and inflammation. The Journal of clinical investigation 125 (2015) 194–207.

134. G. Liu, Y. Bi, L. Xue, Y. Zhang, H. Yang, X. Chen, Y. Lu, Z. Zhang, H. Liu, X. Wang, R. Wang, Y. Chu, and R. Yang, Dendritic cell SIRT1-HIF1alpha axis programs the differentiation of CD4+ T cells through IL-12 and TGF-beta1. Proceedings of the National Academy of Sciences of the United States of America 112 (2015) E957–65.

135. O.U. Kawalekar, R.S. O’Connor, J.A. Fraietta, L. Guo, S.E. McGettigan, A.D. Posey, Jr., P.R. Patel, S. Guedan, J. Scholler, B. Keith, N.W. Snyder, I.A. Blair, M.C. Milone, and C.H. June, Distinct Signaling of Coreceptors Regulates Specific Metabolism Pathways and Impacts Memory Development in CAR T Cells. Immunity 44 (2016) 380–90.

136. L. Liu, Y. Lu, J. Martinez, Y. Bi, G. Lian, T. Wang, S. Milasta, J. Wang, M. Yang, G. Liu, D.R. Green, and R. Wang, Proinflammatory signal suppresses proliferation and shifts macrophage metabolism from Myc-dependent to HIF1alpha-dependent. Proceedings of the National Academy of Sciences of the United States of America 113 (2016) 1564–9.

137. M. Sukumar, J. Liu, G.U. Mehta, S.J. Patel, R. Roychoudhuri, J.G. Crompton, C.A. Klebanoff, Y. Ji, P. Li, Z. Yu, G.D. Whitehill, D. Clever, R.L. Eil, D.C. Palmer, S. Mitra, M. Rao, K. Keyvanfar, D.S. Schrump, E. Wang, F.M. Marincola, L. Gattinoni, W.J. Leonard, P. Muranski, T. Finkel, and N.P. Restifo, Mitochondrial Membrane Potential Identifies Cells with Enhanced Stemness for Cellular Therapy. Cell metabolism 23 (2016) 63–76.

138. L. Fazekas, F. Horkay, V. Kekesi, E. Huszar, E. Barat, R. Fazekas, T. Szabo, A. Juhasz-Nagy, and A. Naszlady, Enhanced accumulation of pericardial fluid adenosine and inosine in patients with coronary artery disease. Life sciences 65 (1999) 1005–12.

139. C.E. Markowitz, S. Spitsin, V. Zimmerman, D. Jacobs, J.K. Udupa, D.C. Hooper, and H. Koprowski, The treatment of multiple sclerosis with inosine. Journal of alternative and complementary medicine 15 (2009) 619–25.

140. S. Spitsin, C.E. Markowitz, V. Zimmerman, H. Koprowski, and D.C. Hooper, Modulation of serum uric acid levels by inosine in patients with multiple sclerosis does not affect blood pressure. Journal of human hypertension 24 (2010) 359–62.

141. S.-P.D.I. Parkinson Study Group, M.A. Schwarzschild, A. Ascherio, M.F. Beal, M.E. Cudkowicz, G.C. Curhan, J.M. Hare, D.C. Hooper, K.D. Kieburtz, E.A. Macklin, D. Oakes, A. Rudolph, I. Shoulson, M.K. Tennis, A.J. Espay, M. Gartner, A. Hung, G. Bwala, R. Lenehan, E. Encarnacion, M. Ainslie, R. Castillo, D. Togasaki, G. Barles, J.H. Friedman, L. Niles, J.H. Carter, M. Murray, C.G. Goetz, J. Jaglin, A. Ahmed, D.S. Russell, C. Cotto, J.L. Goudreau, D. Russell, S.A. Parashos, P. Ede, M.H. Saint-Hilaire, C.A. Thomas, R. James, M.A. Stacy, J. Johnson, L. Gauger, J. Antonelle de Marcaida, S. Thurlow, S.H. Isaacson, L. Carvajal, J. Rao, M. Cook, C. Hope-Porche, L. McClurg, D.L. Grasso, R. Logan, C. Orme, T. Ross, A.F. Brocht, R. Constantinescu, S. Sharma, C. Venuto, J. Weber, and K. Eaton, Inosine to increase serum and cerebrospinal fluid urate in Parkinson disease: a randomized clinical trial. JAMA neurology 71 (2014) 141–50.

142. Y. Lou, G. Wang, G. Lizee, G.J. Kim, S.E. Finkelstein, C. Feng, N.P. Restifo, and P. Hwu, Dendritic cells strongly boost the antitumor activity of adoptively transferred T cells in vivo. Cancer research 64 (2004) 6783–90.

143. T.W.-M. Fan, Sample Preparation for Metabolomics Investigation. in: T.W.-M. Fan, A.N. Lane, and R.M. Higashi, (Eds.), The Handbook of Metabolomics: Pathway and Flux Analysis, Methods in Pharmacology and Toxicology. DOI 10.1007/978-1-61779-618-0_11, Springer Science, New York, 2012, pp. 7–27.

144. T.W. Fan, M.O. Warmoes, Q. Sun, H. Song, J. Turchan-Cholewo, J.T. Martin, A. Mahan, R.M. Higashi, and A.N. Lane, Distinctly perturbed metabolic networks underlie differential tumor tissue damages induced by immune modulator beta-glucan in a two-case ex vivo non-small-cell lung cancer study. Cold Spring Harb Mol Case Stud 2 (2016) a000893.

145. H.N. Moseley, Correcting for the effects of natural abundance in stable isotope resolved metabolomics experiments involving ultra-high resolution mass spectrometry. BMC bioinformatics 11 (2010) 139.

146. A.N. Lane, T.W. Fan, and R.M. Higashi, Isotopomer-based metabolomic analysis by NMR and mass spectrometry. Methods Cell Biol 84 (2008) 541–88.

147. T.W.-M. Fan, and A.N. Lane, Structure-based profiling of Metabolites and Isotopomers by NMR. Progress in NMR Spectroscopy 52 (2008) 69–117.

148. A. Spandidos, X. Wang, H. Wang, and B. Seed, PrimerBank: a resource of human and mouse PCR primer pairs for gene expression detection and quantification. Nucleic Acids Res 38 (2010) D792–9.

149. R. Wang, G. He, M. Nelman-Gonzalez, C.L. Ashorn, G.E. Gallick, P.T. Stukenberg, M.W. Kirschner, and J. Kuang, Regulation of Cdc25C by ERK-MAP kinases during the G2/M transition. Cell 128 (2007) 1119–32.

150. A. Moon, and W.J. Rhead, Complementation analysis of fatty acid oxidation disorders. The Journal of clinical investigation 79 (1987) 59–64.

151. M. Buzzai, D.E. Bauer, R.G. Jones, R.J. Deberardinis, G. Hatzivassiliou, R.L. Elstrom, and C.B. Thompson, The glucose dependence of Akt-transformed cells can be reversed by pharmacologic activation of fatty acid beta-oxidation. Oncogene 24 (2005) 4165–73.

152. K. Brand, J.F. Williams, and M.J. Weidemann, Glucose and glutamine metabolism in rat thymocytes. Biochem J 221 (1984) 471–5.

