## Supplementary material for "Inosine is an alternative carbon supply that supports effector T cell proliferation and antitumor function under glucose restriction": Table S1

| Antibody name | Cat# | Vendor |
|---|---|---|
| Mouse anti-CD3 mAb | BE0001-1 | BioXcell |
| Mouse anti-CD28 mAb | BE0015-1 | BioXcell |
| Anti mouse CD8-APC-Cy7 | 100714 | Biolegend |
| Anti human/mouse Granzyme B-FITC | 515403 | biolegend |
| Anti mouse TNF-α- APC | 17-7321-82 | eBioscience |
| Anti mouse IFN-γ- PE-Cy7 | 25-7319-82 | ebioscience |
| Anti-human CD8a Antibody - APC-Cy7 | 300926 | Biolegend |
| Anti-human TNF-α -PE | 502909 | Biolegend |
| Anti-human IFN-γ- PE-Cy7 | 506518 | Biolegend |
| Anti mouse CD45.2 - PerCP-Cyanine5.5 | 45-0454-82 | eBioscience |
| Anti mouse CD3-FITC | 11-0031-85 | eBioscience |
| InVivoMAb anti m PD-L1 | BE0101 | BioXcell |
| Anti-human CD3 (OKT-3) | BE0001 | BioXcell |
| Anti-human/monkey CD28.2 | BE0291 | BioXcell |
| anti-PNP | sc-271890 | Santa Cruz |
| anti-actin | sc47778 | Santa Cruz |

| Reagents name | Cat#/Vendor |
|---|---|
| Recombinant human IL-2 | 200-02, Peprotech |
| Recombinant murine IL-2 | 212-12, Peprotech |
| Cell Stimulation Cocktail (plus protein transport inhibitors) (500X) | 00-4975-93,eBioscience |
| carboxyfluorescein diacetate succinimidyl ester(CFSE) | Invitrogen |
| 7-amino-actinomycin D(7AAD) | 420404,Biolegend |
| D-(+)-Glucose | G7021- sigma |
| IncuCyte™ Caspase-3/7 Apoptosis Reagent | Essen BioScience 4440 |
| Inosine | I4125-sigma |
| Adenosine | 164040050, ACROS |
| D-(+)-Glucose | G7021, sigma |
| RPMI 1640 Medium, No Glucose | 11-879-020, Gibco |
| Retronectin | T100, Takara/clontech |
| Human gp100 | RP20344, GenScript |
| Forodesine Hydrocholoride | HY-16209-MedChem Express |
| $^{13}C_6$-Glucose | CLM-1396-Cambridge Isotope Lab |
| $^{13}C_5$-Inosine | NUC-072-OMICRON |
| [$^{14}C_5$]-glutamine | ART0115, American Radiolabeled Chemicals |
| [9,10-3H]-palmitic acid | MT 845, Moravek |
